# LSD Reconfigures Cortical Dynamics Through Faster Brain Rhythms and Increased Fractal Dimension

**DOI:** 10.64898/2026.01.28.702361

**Authors:** Venkatesh Subramani, Timothy Nest, Annalisa Pascarella, Jérémy Brunel, Yorguin José Mantilla Ramos, Yann Harel, Suresh Muthukumaraswamy, Robin Carhart-Harris, Giulia Lioi, Nicolas Farrugia, Karim Jerbi

## Abstract

Lysergic acid diethylamide (LSD) profoundly alters conscious experience, yet the electrophysiological mechanisms by which it reshapes neural dynamics remain incompletely understood. A hallmark of psychedelic states is widespread cortical desynchronization, typically inferred from reductions in spectral power, but whether such effects reflect genuine weakening of neural oscillations or are confounded by shifts in oscillatory peak frequencies remains unresolved. Here, we address this gap by combining source-resolved magnetoencephalography (MEG), spectral parameterization, temporal complexity metrics, and interpretable machine learning in an LSD versus placebo design, with and without music. We show that LSD induces robust, spatially structured increases in alpha and beta peak frequencies alongside genuine attenuation of oscillatory power, with these effects displaying partly dissociable cortical patterns. Beyond rhythmic activity, LSD is associated with flattening of the aperiodic 1/f spectral slope and increased neural signal fractality and complexity, preferentially affecting sensory, language, emotion, and imagery-related networks while sparing motor cortex. Machine-learning analyses further identify peak-frequency shifts, aperiodic parameters, and complexity measures as key discriminators of the psychedelic state. Music does not robustly amplify these neural signatures and instead shows a trend toward attenuation. Together, these findings provide a comprehensive electrophysiological account of how LSD reorganizes large-scale human brain dynamics and highlight features that may differentiate its neural signature from that of other psychedelics.

## 1 Introduction

Psychedelics are potent psychoactive compounds that elicit profound alterations in conscious experience, spanning heightened sensory perception, vivid imagery, and distortions in the sense of self [1]. Beyond their characteristic phenomenology, these compounds offer a powerful experimental lever for perturbing large-scale brain dynamics and probing the neural mechanisms through which the brain gives rise to mind [2, 3, 4, 5]. Psychedelics have been used in rituals and cultural settings for thousands of years [6], often alongside music [7]. Classic psychedelics—including psilocybin, *N, N* -dimethyltryptamine (DMT), and lysergic acid diethylamide (LSD)—exert their profound effects primarily through agonism at the 5-HT2*A* receptor. Among these, LSD is uniquely characterized by its additional affinity for dopaminergic receptors [8], its extreme potency (producing effects at remarkably low doses), and its extended receptor kinetics. Despite these well-known pharmacological properties, the precise manner in which LSD perturbs cortical dynamics—spanning oscillatory rhythms, aperiodic spectral structure, and temporal signal complexity—and how these effects are shaped by sensory stimuli such as music, remains poorly understood.

Modern neuroimaging has played a central role in uncovering how psychedelics alter large-scale brain function and core aspects of conscious experience, a process thought to support their therapeutic effect [5, 9, 10]. These mechanisms have been investigated across different modalities, including positron emission tomography (PET) [11], functional MRI (fMRI) [12], electroencaphelography (EEG) [13], and magnetoencephalography (MEG) [14]. fMRI has been especially influential in understanding the psychedelic state (e.g., [15]). However, the fMRI signal reflects neural activity only indirectly via blood oxygenation, and psychedelics strongly modulate neurovascular processes [16]. As a result, psychedelic-induced BOLD changes may partly reflect disruption in neurovascular coupling rather than neural dynamics per se [16]. Although simultaneous EEG–fMRI studies are beginning to address these issues [17], direct electrophysiological measurements remain essential for capturing the fast-changing dynamics that characterize the psychedelic state.

M/EEG provides a direct measurement of the brain’s electrophysiological activity at a millisecond scale, allowing precise characterization of oscillatory dynamics [18, 19]. Early electrophysiological studies revealed a robust broadband desynchronization under psychedelics—including psilocybin [14], LSD [12], and DMT [13]. However, spectral power of the neural signals reflects a mixture of periodic (oscillatory) and aperiodic (1/*f* -like) components [20, 21], and changes in one component can be misattributed to the other, leading to ambiguous interpretation of spectral power. This has motivated a shift toward explicit spectral parameterization to separately quantify rhythmic and arrhythmic activity [22, 13, 23]. While broadband desynchronization appears as a robust, cross-psychedelic effect [14, 13, 12], emerging evidence suggests that LSD may additionally modulate spectral peak frequencies. Carhart-Harris and colleagues [12] reported subtle shifts in the alpha peak in posterior MEG sensors, a modulation not reported for DMT [13] and only inconsistently for psilocybin [23]. These observations raise the possibility that frequency-specific perturbations, particularly within the alpha and beta band, constitute a distinguishing signature of LSD relative to other serotonergic psychedelics. Yet, the existing studies have not mapped how these peak-frequency shifts are organized across the cortex, leaving the spatial expression of oscillatory power reduction as well as alpha and beta peak modulation an open question in psychedelic neuroscience. Importantly, traditional analyses using fixed canonical bands can produce misleading results when oscillatory peaks shift. Because the power spectrum follows a 1/*f* -like decay, a peak that shifts upward may appear to lose strength simply because it is measured against a lower baseline, erroneously suggesting a desynchronization when the oscillation has merely changed frequency. To our knowledge, no previous study has analyzed LSD-induced spectral power changes while taking the peak shifts into account.

Music is a powerful sensory stimulus that shapes perception, emotion, and internal experience [24, 25]. Historically and across cultures, it has accompanied ritual practices and psychedelic experiences, suggesting a close relationship between music and altered states of consciousness. The heightened sensory perception and mood characterizing the psychedelic state synergizes with, and potentiates, music’s evocative power. Music therefore serves as a potent contextual factor that shapes internal experience and provides a useful framework for investigating altered brain dynamics [26, 27]. Set (Mindset) and Setting (surrounding factors) play a crucial role in a psychedelic trip [28]. Music has emerged as a particularly influential setting variable and its usefulness in psychedelic-assisted therapy is now fully recognized [29, 30]. Empirical studies have shown that music under psychedelics increases emotional response [31], enhances mental imagery [32], supports meaning-making [33]. In addition, converging evidence suggests that emotional breakthrough and psychological insight may be more robust and clinically predictive dimensions of psychedelic experience [34, 35, 36, 37, 38, 39]. These constructs arguably capture tangible processes, such as affective release and cognitive restructuring, and may therefore offer a more direct bridge between phenomenology and underlying neurobiology. Despite this extensive phenomenological and therapeutic work, the neural mechanisms underlying these effects remain poorly understood. While M/EEG studies have shown attenuated top-down suppression of prediction error under LSD during basic auditory processing [40]— and increases in the richness of the neural signal during music listening [41], a mechanistic account of how music modulates the oscillatory and temporal dynamics of the psychedelic brain is yet to be established.

Beyond their effects on spectral structure, psychedelics are known to alter the temporal complexity of neural activity. Lempel-Ziv complexity (LZC), a prominent metric in psychedelic research, quantifies the diversity of neural signals by measuring the rate of appearance of unique temporal patterns, effectively indexing the richness of the time series. Numerous studies consistently reported an increased LZC under psilocybin, LSD, and DMT [23, 41, 13, 42]. In contrast, measures of fractality such as the Higuchi fractal dimension (HFD) quantify the degree to which brain dynamics are self-similar across time scales, capturing scale-free temporal structure. It therefore complements LZC by revealing how complexity is organized across scales rather than how novel the signal is over time. HFD has been applied in other altered states of consciousness [43], but has received far less attention in psychedelic research. Recent work suggests that psychedelic-induced rise in complexity can be modulated by external context, such as whether one attends to a stimulus or not [41]. This raises another question we address in the present study: to what extent are LSD-induced changes in temporal complexity intrinsic to the drug state, and to what extent are they shaped by structured sensory input such as music?

Machine learning (ML) offers a data-driven framework for distinguishing brain states and has been successfully applied to EEG and MEG data across a range of altered states, including anaesthesia [44], meditation [43], sleep modulation [45], and psychedelic drug effects [46, 47]. Beyond classification accuracy, ML models can provide insight into the neural features that drive drug state differences, particularly through feature-importance analyses. Such approaches offer a complementary perspective to traditional group-level statistics by highlighting neurophysiologically meaningful variables that contribute most strongly to discriminability. In this study, we leverage interpretable ML to characterize how LSD — and its interaction with music — manifests in large-scale cortical dynamics.

In this study, we ask three questions: (Q1) To what extent does LSD produce region-specific modulations of the peak frequency of neural oscillations within the alpha and beta band? (Q2) To what extent does LSD modulate the power of alpha and beta oscillations once the confounding effects of peak frequency shifts are accounted for? (Q3) Does music modulate LSD-induced alterations in neural temporal organization, including oscillatory dynamics and signal complexity? To address these questions, we apply spectral parameterization, oscillatory peak–frequency analysis, and temporal-complexity metrics (LZC and HFD) to source-reconstructed MEG data collected under LSD and placebo (PLA), with and without music. We further use machine-learning models to quantify the discriminative value of these features. Based on prior work and the mechanistic considerations outlined above, we formulated three hypotheses: (H1) LSD induces region-specific shifts in oscillatory peak frequencies; (H2) LSD induces genuine attenuation of oscillatory power independent of peak-frequency shifts; (H3) music potentiates LSD-induced neural effects by amplifying changes in oscillatory organization and temporal complexity.

## 2 Materials and Methods

### 2.1 Data

In this study, we analyzed MEG data from 17 participants collected by Carhart-Harris and colleagues [12]. The experimental setup followed a within-subject single-blind design, with the order counterbalanced. MEG recordings took place on two different days during which participants under the influence of either intravenous LSD (75*µ*g) or Placebo (saline), rested, or performed passive tasks. In this study, we used a subset of the conditions during which the participants either had their eyes-closed (NoMusic condition), or eyes-closed with music (Music condition). The duration of each recording was 7 minutes. For the music condition, participants listened to two seven-minute long excerpts from the electro-acoustic ambient album *Yearning* by Robert Rich and Lisa Moskow. LSD and PLA sessions were 14 days apart to prevent drug cross-over effect. Details about the criteria for the recruitment of the participants are outlined in the original data study [12]. Approximately 4 hours post-injection MEG was recorded using a CTF-275 system at 1200 Hz (0-300 Hz bandpass), with four excessively noisy sensors turned off. Following each MEG run, participants completed in-scanner subjective rating scales assessing several phenomenological dimensions such as mood and imagery (see supplementary Fig. S5 for a complete list and raw scores).

### 2.2 Preprocessing

Our preprocessing pipeline resembles the approach used in the original study, but incorporates additional automated procedures to improve artefact removal implemented using the Autoreject package [48] (Figure 1). The pipeline consists of four steps: 1) filtering, visual inspection, and epoching; 2) first pass of Autoreject; 3) ICA-based artefactual components removal; 4) second pass of Autoreject. In particular, the continuous MEG recordings were bandpass filtered between 1 and 120Hz (implemented in MNE v1.8’s Raw.filter), and notch-filtered at 50 and 100 Hz (implemented in MNE’s Raw.notch_filter). As with the original study, we segmented the continuous MEG signal into 2s epochs (210 epochs per run) [12]. This enabled us to isolate and systematically remove segments containing artefacts. Large, clearly identifiable artefacts were removed through visual inspection. The majority of the epochs passed our visual inspection (Mean epochs per subject: 205.5; SD : 9.3), and epoch retention was rather consistent across the LSD and PLA groups (paired t-test; t=-1.61; *p <* 0.12). The first pass of Autoreject was run in *mask-only* mode, i.e. Autoreject was employed to identify and label the bad epochs, which were then excluded prior to ICA. ICA decomposition (Picard algorithm, see [49]) enabled us to identify and exclude the artefactual components. The remaining components were back-projected to reconstruct the signal across all epochs, including those initially marked as bad or interpolated. To further attenuate residual artefacts, a second pass of Autoreject was applied to the reconstructed data. Epochs labeled as good or interpolated after this step were retained as clean data. The final cleaned dataset contained approximately 200 epochs per run on average (SD = 20), again with no significant difference in epoch retention between LSD and placebo conditions (*t* = −1.69, *p <* 0.12).

**Figure 1:**
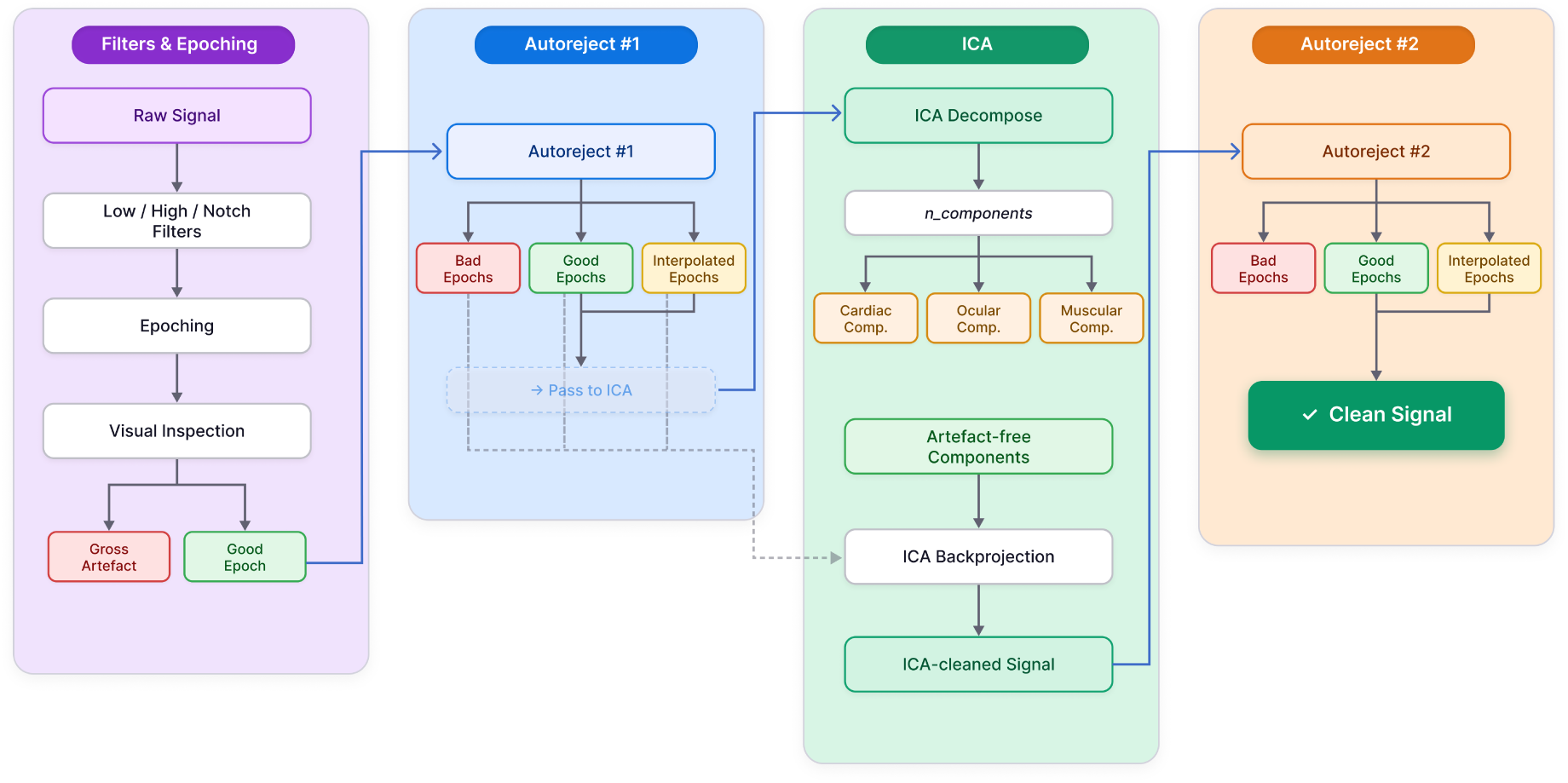
Overview of the preprocessing pipeline. The pipeline begins with filtering, visual inspection, and segmentation of the continuous MEG signal into 2-s epochs, following the procedure of Carhart-Harris et al. [12]. Subsequent cleaning is performed using a quasi-automated, three-stage workflow. In the first stage, Autoreject is applied in *mask-only* mode to identify bad epochs while preserving the full dataset for later reconstruction. The good and interpolated epochs are then subjected to ICA decomposition. Components reflecting cardiac, ocular, or muscular artefacts are removed, and the remaining components are back-projected onto all epochs (good, interpolated, and previously flagged), producing an “ICA-cleaned” signal. Because residual artefacts may persist after ICA back-projection, a second Autoreject pass is performed to reject any remaining contaminated epochs, yielding the final cleaned dataset.

### 2.3 Source Localization

Structural T1-weighted MRI scans were available for each participant in both LSD and PLA sessions [12]. These anatomical images were processed using FreeSurfer’s standard reconstruction pipeline (recon-all, v7.4.1), which performs motion correction, intensity normalization, skull stripping, and white–gray matter surface segmentation. All reconstructions were visually inspected for accuracy prior to source analysis. These subject-specific cortical surfaces were used to estimate the cortical sources for the MEG signal. Source localization requires both a forward model and an inverse solver. For the forward model, we generated Boundary Element Model (BEM) surfaces using FreeSurfer’s watershed algorithm (implemented in MNE). Because MEG is relatively insensitive to conductivity differences across head tissues [19], we employed the standard single-layer (inner-skull, conductivity = 0.3) BEM model commonly used in MEG source estimation. Individual source spaces were defined on the ico5 spacing (20484 vertices). MEG sensor positions were co-registered to anatomy using three fiducials (nasion, left and right preauricular points), yielding a 4×4 affine transformation matrix mapping MEG coordinates to each participant’s MRI space. The inverse solution requires specification of the noise covariance matrix. Because empty-room noise recordings were not available for this dataset, we adopted the commonly accepted alternative of using an identity covariance matrix, which assumes uniform noise across sensors. We set the orientation constraints to be mostly perpendicular to the cortex (MNE’s loose = 0.2). Source estimation was performed using the dynamic Statistical Parametric Mapping (dSPM) inverse operator [50], yielding time series at each cortical vertex for each epoch.

For group-level analyses, individual source estimates were morphed (source space morphing) to the FreeSurfer fsaverage (v5) template. The morphed source time courses were then parcellated onto the HCP-MMP atlas [51], implemented in MNE’s extract_label_time_course. In particular, we extracted the ROI’s activity by averaging over the corresponding fsaverage vertices, resulting in N = 360 regional time series per participant per condition. These source-projected signals were used to compute temporal metrics such as LZC and HFD. Power spectral density (PSD) estimates were obtained at the source level (20484 vertices) using the multitaper method implemented in MNE’s compute_psd_epochs, followed by morphing and parcellation to yield PSDs for each cortical region in the HCP-MMP atlas.

### 2.4 Spectral Parametrization

Electrophysiological signals contain both periodic (oscillatory) and aperiodic (1/*f* -like) components [20]. Because these components contribute jointly to the power spectrum, analyzing raw, non-parametrized spectra can conflate the interpretation. To disentangle these contributions, we parameterize each power spectrum using the Fitting Oscillations and One-Over-f toolbox [20]. To this end, we fit spectral parametrization models on the epochs-averaged spectra with a frequency range of 1 - 120 Hz using the *fixed* aperiodic mode, with max number of peaks set to 5. Prior to fitting, we applied a narrow one-dimensional interpolation around the powerline frequencies and the harmonics (50 Hz and 100 Hz; 3-Hz notch width) to prevent sharp notch artefacts from distorting the parametrization. Model performance and goodness-of-fit metrics are reported in the Supplementary Materials (Supplementary Figure S4). The spectral parametrization model decomposes each spectrum into a set of Gaussian peaks representing oscillatory activity and an aperiodic component, characterized by the offset and the slope which corresponds to the exponent of the 1/*f* -like distribution of the power spectrum. We extracted these parameters and tested for the perturbations induced by LSD and by LSD x music interaction. In addition, we examined the pure oscillatory component and the band peaks as identified by the spectral parametrization model. The oscillatory activity was robustly present in alpha and beta bands, and notably present in high-gamma band (Supplementary Figure S1). Consequently, in this study, we focused on alpha, beta and high-gamma activity. We computed the oscillatory power (without 1/*f* component) in the canonical frequency range (results presented in Supplementary Figure S6). However, we observed shifts in the alpha and beta peaks that may confound the band power analysis. Thus, as spectral feature we considered also the Peak Power, defined as the maximum pure oscillatory activity in the specified band. Also, to assess the robustness of our parametrization pipeline, we also performed the analysis by fitting the model for a narrower range of 1 - 45 Hz. The corresponding results can be found in the supplementary material Supplementary Figure S4.

### 2.5 Complexity

Lempel–Ziv complexity quantifies the compressibility of the EEG or MEG signals, i.e. how diverse the patterns are in the signal [52]. While it is not an entropy measure per se, increases in LZC are commonly interpreted as reflecting more diverse and less constrained neural dynamics, often associated with higher entropic brain states [53]. Computing LZC involves two steps: First, the continuous time series is binarized, typically using the mean (or median) as a threshold such that values greater than the threshold are assigned “1” and all others “0”. Second, the number of distinct patterns in the resulting binary sequence is counted, yielding the LZC value. Lower LZC indicates more regular, predictable activity, whereas higher LZC reflects greater temporal diversity. LZC has been successfully applied to electrophysiological data, especially with the altered states of consciousness, including the recent work investigating the effects of external stimulation under LSD [41]. Although Mediano, Rosas et al. [41] examined the effects of LSD on brain complexity across multiple task contexts, we extend this work in two important ways. First, we specifically quantify LZC in the context of interaction between LSD and music, directly probing how sensory context modulates signal richness under psychedelics. Second, we compute LZC using our own preprocessing pipeline, which is appreciated from a replicability standpoint. In this work, we computed LZC for a) broadband signal (1-120Hz, main text); b) bandlimited signals within each canonical frequency band (alpha through high-gamma; supplementary).

LZC was computed by concatenating all artefact-free epochs. The binarization threshold was set to the average of the signal. Because the duration of the concatenated signals varied slightly across participants after preprocessing, we normalized [54] the LZC by 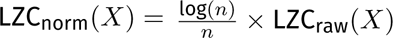, where X denotes the signal, and n its length.

### 2.6 Fractality

Fractal dimension quantifies scale-invariant temporal structure in neural signals. Here, we estimate fractal dimension using HFD [55], which captures self-similarity across time scales in EEG and MEG data, capturing scale-free temporal structure [56, 57]. It complements Lempel–Ziv complexity by characterizing how signal complexity is organized across temporal scales, rather than how diverse the signal patterns are over time. The HFD algorithm estimates how the “length” of a signal changes as a function of the scale at which the signal is sampled. HFD computation involves a 4-step process. First, for each integer *k* = 1*, ߪ, K*_max_, the original time series *X*(1)*, X*(2)*, ߪ, X*(*N*) is used to construct *k* subsampled sequences

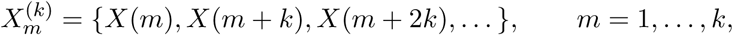

each beginning at offset *m* and sampled at interval *k*. Second, for each sequence 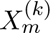, a curve length *L_m_*(*k*) is computed using the normalized sum of absolute successive differences, and the mean curve length across all offsets is

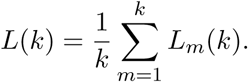

Third, the set of values {*L*(*k*)} is then examined in log–log space, where fractal signals exhibit a systematic decrease in *L*(*k*) as *k* increases. Finally, the fractal dimension is obtained as the slope *D* of the linear regression of log *L*(*k*) versus log(1/*k*). HFD ranges between 1 and 2, increasing asymptotically and saturates at some 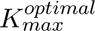, a hyperparameter (e.g., [58]).

We computed the HFD across ROIs, subjects, drug and task conditions as a function of *K*_max_ ∈ [2, 50]. To select the 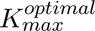, we examined the HFD-*K*_max_ curve and identified the saddle point by applying a knee-detection algorithm. The resulting knee index yielded an optimal value of 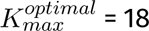 = 18, which we adopted for all the HFD computations. We computed the HFDs for each epoch, and subsequently averaged over epochs within each recording. All fractal-dimension estimates were obtained using the higuchi_fd function from the antropy Python package.

### 2.7 Functional Decoding

The LSD-induced neurophysiological changes were mapped onto the large-scale cognitive and affective systems using Functional Decoding, with NiMare [59], following previous studies [60, 61, 62]. The aim of this analysis is to identify which cognitive or affective processes are most strongly associated with the spatial distribution of LSD-induced effects. Functional decoding relies on the Neurosynth database, a large-scale, automated meta-analytic resource that aggregates activation coordinates from more than 14,000 neuroimaging studies (over 500,000 statistical maps). Neurosynth organizes the literature into topics [63], which are latent semantic components derived from the co-occurrence of terms in article abstracts and their associated activation patterns. Each topic therefore represents a broad cognitive or affective construct (e.g., visual perception, autobiographical memory, emotion, language) rather than a single experimental task or region. For each electrophysiological feature, the unthresholded statistical contrast map (LSD and PLA) was rank-ordered and segmented into 10 equally sized bins (10% increments). Each decile mask was binarized and submitted to the ROIAssociationDecoder in NiMARE, which computes the spatial correspondence between the input masks and meta-analytic activation maps (see [63] for details). The decoder returns correlation coefficients reflecting the degree to which that spatial pattern is associated with each Neurosynth topic. These correlations were transformed into z-statistics, and only associations surviving a threshold of *p <* 0.001 were retained. Following [60], topic terms were ordered based on the weighted average of each topic’s z-scores across the ten deciles. This ranking highlights the cognitive and affective domains that best characterize the regions showing the strongest LSD-induced electrophysiological changes.

### 2.8 Phenomenology Regression analysis

We investigated how LSD-induced neurophysiological changes relate to participants’ subjective experience by regressing the changes in the neurophysiological features against the subjective ratings. We quantified the relation between the two variables using Spearman’s correlation *ρ*, a non-parametric rank-based test that capture their monotonic relationship. The resulting strength and relation are compressed into the correlation coefficient *ρ*, that ranges between -1 and 1 indicating negative and positive relation between the two variables. Analyses were conducted at two spatial scales: i) global scale (i.e., region-averaged); ii) region-resolved scale, which enables detection of localized brain–experience relationships that would be obscured in global averages. Given the limited sample size and multiple comparisons inherent to region-resolved analyses, these regressions should be interpreted as exploratory. The corresponding results can be found in the supplementary figures S2 and S3.

### 2.9 Statistics and Reproducibility

For each feature, we contrasted LSD and PLA by applying paired statistical tests with multiple comparison correction. In particular, for the global spectral analyses (see 2A), we performed non-parametric cluster-level tests (implemented in MNE’s permutation_cluster_1samp_test), in which the temporal adjacency was defined along the frequency points (Hz). Clusters were formed by grouping consecutive frequency bins exhibiting similar test-statistic values (both in sign and magnitude), using a cluster-forming threshold corresponding to a *p <* 0.05 (see [64] for details).

For all other features, we employed non-parametric permutation-corrected tests [64] (implemented in MNE’s permutation_t_test). Empirical test statistics were computed from paired LSD-PLA contrasts, and evaluated against a null distribution obtained by permuting the regions (N=50000 permutations). For each region, the p-value corresponded to the percentile rank of the observed statistic *T_obs_*within this surrogate distribution, following the procedure outlined in [64]. Statistical significance was defined at *p <* 0.05. In cases where the LSD-induced perturbations were broad and spatially extensive throughout the cortex, we additionally highlighted regions showing the strongest effects by marking the strongest decile of the resulting statistical distribution (e.g., t-values).

For the regression analyses linking neural features to subjective experience, we controlled for multiple comparisons using false discovery rate (FDR [65]) correction at *p <* 0.05. We show in the supplementary material for the comparisons (Supp. Fig. S2 and S3).

In other scenarios such as the interaction effect of LSD and music, we report that the effects are nominally significant (i.e. *p <* 0.05 uncorrected), reflected in all features.

The *Main Effect Drug* quantifies the effect of LSD regardless of the task condition, whereas *Main Effect Music* captures the effect of Music irrespective of the drug state. The *Interaction effect* (LSD x Music) tests whether music modulates the way LSD affects the brain. Finally, to further decompose the LSD x Music interaction, we looked at the Drug effect under music (*Simple Effect Music*) and without music (*Simple Effect NoMusic*) to elucidate how the LSD x Music interaction manifests.

### 2.10 Machine Learning

We used Random Forest classifier, an ensemble tree-based ML model to decode LSD versus PLA states from the range of temporal and spectral features (oscillatory power, peak frequencies, LZC, and HFD). We chose a Random Forest classifier as it can capture potential non-linear relationships and interactions across heterogeneous feature domains, and because it generally provides robust performance without requiring assumptions about linear separability or feature independence. We extract the features that are important to discriminate between the two states, termed Feature Importances. We conducted two complementary analyses by building single-domain and multi-domain models.

In the single-domain Model, each classifier operated on one feature family at a time (e.g., alpha Peak Power, beta peak frequency, exponent, LZC), using the set of regional values as model inputs. In this formulation, the ML “features” correspond to cortical regions, and the resulting Feature Importances identify the regions whose values most strongly discriminate LSD from PLA. This notion is analogous to the classical statistics, where the region is significantly different when the LSD and PLA distributions are not invariant. In contrast, the multi-domain model pooled all temporal and spectral features across all regions into a single classifier, enabling us to determine which feature families, and which regions within them, carry the greatest predictive weight *overall*. Such multi-domain decoding models are increasingly used to investigate altered states of consciousness—including those induced by caffeine during sleep [45] and meditation [66]—and offer a complementary, more holistic perspective on which neural markers most reliably distinguish drug states.

Training procedures were identical for both single-domain and multi-domain models. Model hyperparameters were optimized via nested StratifiedGroupKFold cross-validation (outer K=4, inner K=3) with grid search, where the inner loop handled hyperparameter selection, keeping outer test folds unseen and yielding an unbiased performance estimate. To assess significance, we constructed surrogate null distributions for both accuracy and Feature Importances by permuting the target labels 1,000 times. For each permutation, the full StratifiedGroupKFold training procedure was repeated, and surrogate accuracies and surrogate Feature Importances were obtained by averaging across folds. Empirical values were then compared against their respective null distributions to obtain permutation-based p-values, which quantify the probability that feature–label associations arise under the assumption of independence. Feature Importances surviving *p <* 0.05 thus reflect permutation-corrected, multiple-comparison–controlled indices of discriminability between LSD and PLA. In addition to reporting accuracy and feature importances in the main text, we also report the AUC-ROC plots in supplementary (Supp. Fig. S9).

## 3 Results

In what follows, we report four complementary sets of findings aligned with our three research questions. First, we characterize how LSD induces region-specific shifts in alpha and beta peak frequencies (Q1). Second, we isolate genuine changes in oscillatory power after accounting for these peak displacements, allowing us to dissociate true dampening from apparent reductions driven by frequency shifts (Q2). Third, to contextualize these oscillatory changes at the systems level, we examine how these neurophysiological effects are organized across large-scale functional systems using meta-analytic functional decoding. Finally, we assess how these neurophysiological effects are modulated by music, addressing the interaction between pharmacological and contextual influences (Q3).

### 3.1 LSD induces genuine attenuation of alpha and beta oscillations

We examined the main effect of LSD on oscillatory power by parameterizing cortical power spectra and isolating the periodic component (Figure 2). At the whole-brain level, power was first averaged across all cortical regions. In the total spectra (Figure 2A, top), LSD was associated with a broad reduction in low-frequency power and a relative increase at higher frequencies; however, these differences did not survive cluster-based multiple-comparison correction (*p <* 0.05). After removing the aperiodic 1/*f* component (Figure 2A, bottom), a clearer pattern emerged: LSD robustly weakened oscillatory activity from theta through beta bands (up to 30 Hz), while mid-gamma power (60–90 Hz) showed a significant increase (*p <* 0.05; cluster-corrected). These region-averaged spectra summarize the global spectral effects of LSD; we next examined their spatial distribution across cortical regions.

**Figure 2:**
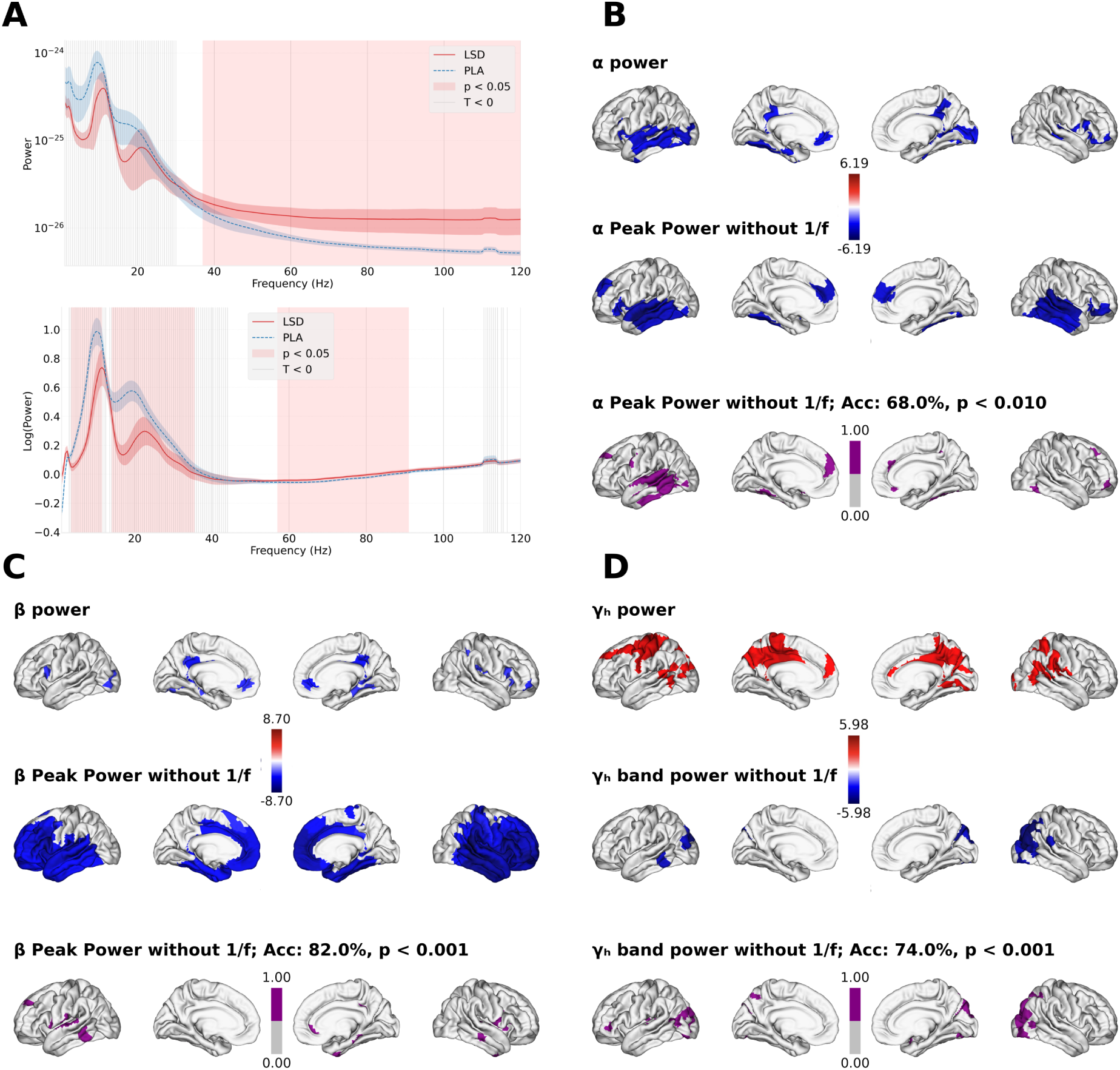
Global and region-resolved spectral effects of LSD (Main Effect Drug). **A** Region-averaged power spectra before spectral parameterization (“raw spectrum”, top) and after removing the aperiodic component (“oscillatory component”, bottom). Gray vertical bands indicate frequency ranges where LSD < PLA (dashed line); red shading indicates statistically significant differences (cluster-corrected, *p <* 0.05). **B-D** Regional effects of LSD on *α* (**B**), *β* (**C**), and high-*γ* (**D**) activity. For each band, the first row shows the contrast maps (permutation-corrected, *p <* 0.05) for power within the canonical frequency band (*α*, *β*, *γ*) when the aperiodic (1/*f* ) component is retained. The second row for *α* and *β* shows contrast maps for power at the oscillatory peak, after isolating periodic activity via spectral parameterization. The contrasts for high-*γ* corresponds to changes in bandlimited oscillatory power. The third row displays permutation-corrected machine-learning feature-importance maps from models decoding LSD vs placebo using Peak Power features. Highlighted regions indicate strong contributions to classification. Decoding accuracies are cross-validated and permutation-corrected.

Using spectral parametrization to probe the presence of genuine oscillatory components, we observed that oscillatory activity is most consistently expressed across cortical regions in the alpha and beta bands, with reliable oscillatory structure also present in the high-gamma range (see Supplementary Figure S1). Accordingly, subsequent analyses focus on these frequency bands. Permutation-based statistical contrasts (LSD vs. PLA; 50,000 permutations) revealed spatially specific effects across the cortex (Figure 2B–D), with negative *t*-values (blue) indicating power suppression and positive values (red) indicating power increases. In the total power spectra (Figure 2B–D, first row), LSD predominantly reduced alpha and beta power while producing focal increases in high-gamma activity. To isolate genuine oscillatory effects, we focused on power at oscillatory peaks (Peak Power), rather than power within canonical frequency bands, as LSD induced systematic shifts in alpha and beta peak frequencies. Peak-based contrasts (permutation-corrected, *p <* 0.05; Figure 2B–C, second row) revealed frequency-specific spatial patterns. Alpha Peak Power reductions were most prominent in temporal regions, whereas beta Peak Power showed widespread suppression across the cortex. In the high-gamma range, oscillatory power was reduced primarily in posterior regions. Across all three bands, analyses no longer showed significant effects in the posterior cingulate cortex, which had appeared prominently in total power contrasts, suggesting that earlier effects were driven by changes in the aperiodic component. Analyses of band-limited power without accounting for peak shifts yielded more spatially extensive suppression (Supplementary Figure S6), indicating that ignoring peak-frequency shifts inflates estimates of drug effects.

To complement inferential statistics, we applied supervised machine-learning to assess the discriminability of LSD and PLA states. Employing machine learning here aims to improve the robustness of our findings, and provide more interpretability to inferential statistics. Random Forest classifiers were trained using regional spectral power features (alpha and beta Peak Power; high-gamma band-limited power) to predict drug condition. Classifiers achieved robust decoding performance (accuracy *>* hl68% Supp. figure S9 for AUC-ROC plots), indicating strong separability between LSD and placebo states. Feature importance analysis revealed that the regions contributing most strongly to classification closely matched those identified by permutation-based *t*-tests (Figure 2B–D, third row). The statistical significance of feature importances was assessed against null distributions generated via label permutation (1,000 iterations), with regions exceeding the 95th percentile deemed significant (*p <* 0.05). The convergence between ML–derived importance maps and classical statistical results across frequency bands supports the robustness and spatial specificity of LSD-induced oscillatory changes.

### 3.2 LSD robustly shifts peaks, flattens slope and increases fractality

Figure 3 summarizes the main effects of LSD on additional spectral and temporal features. Figure 3A shows paired permutation-based statistical contrasts (LSD vs. PLA; *p <* 0.05, corrected; 50,000 permutations), and Figure 3B shows feature-importance maps derived from Random Forest classifiers trained separately on each feature to decode drug condition. Statistical significance of feature importances was assessed using label permutation (*N* = 1,000, *p <* 0.05).

**Figure 3:**
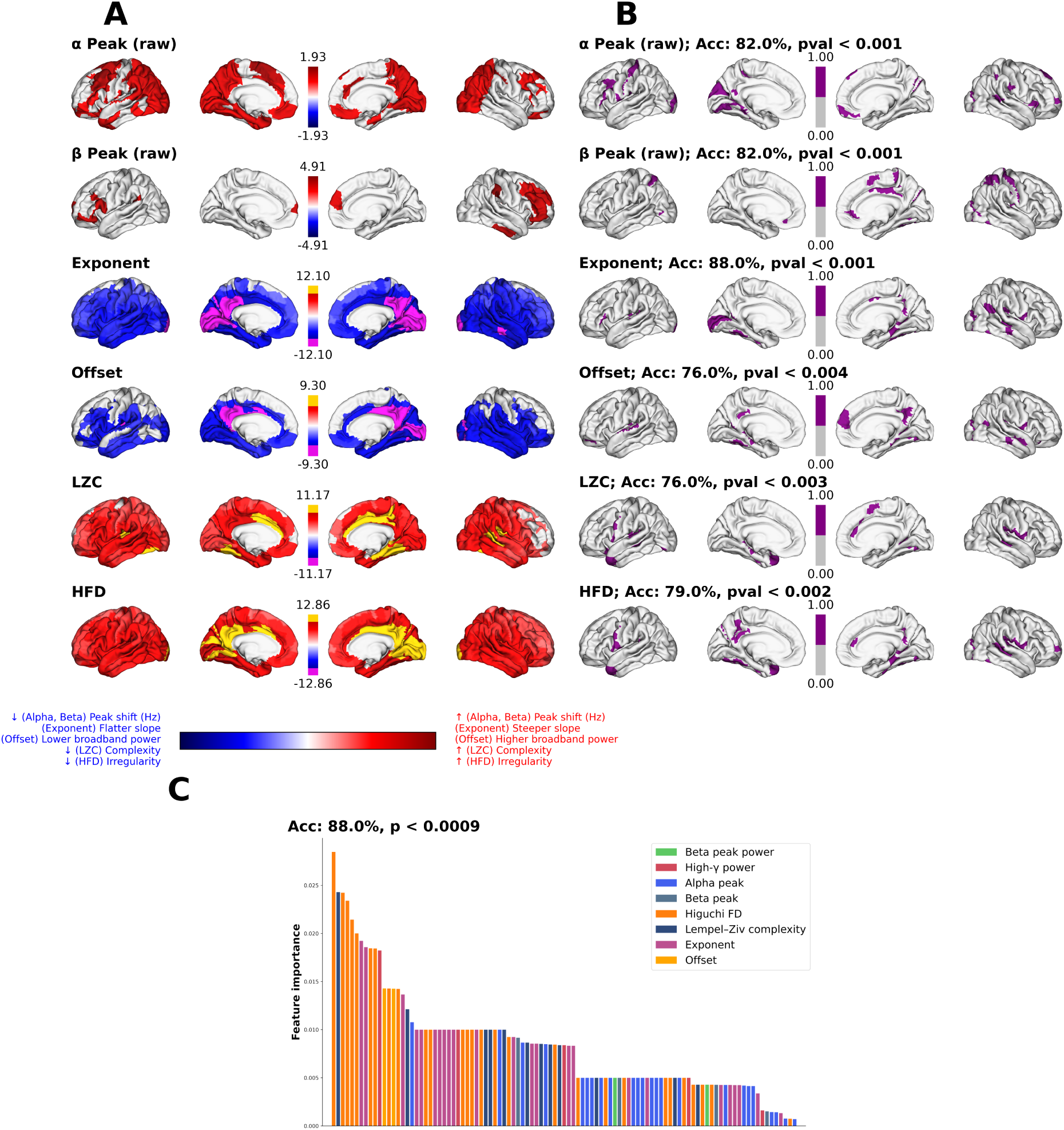
LSD-induced alterations in temporal and spectral electrophysiological features. **A** Region-resolved LSD–PLA contrasts (permutation-corrected) for six features: grand-averaged LSD–PLA differences in *α* and *β* peak frequencies, each projected onto their respective significance masks; aperiodic exponent, aperiodic offset, Lempel–Ziv complexity (LZC), and Higuchi fractal dimension (HFD). For exponent, offset, LZC, and HFD, the color maps reflect t-statistics, and the strongest 10% of the regions are highlighted in yellow and magenta. The corresponding *p* value for the highlighted regions are by the colorbar (*** *p <* 10*^−^*^3^, ***** *p <* 10*^−^*^5^). **B** Machine-learning feature-importance maps (binarized) derived from Random Forest decoders trained to distinguish LSD from PLA based on each feature class separately. Significant regions are determined via permutation-testing of feature importances (1,000 null-label permutations; *p <* 0.05). Classification accuracy and permutation-corrected p-values are reported for each feature. **C** Multi-domain decoding analysis using all feature families together. Bars represent permutation-corrected feature importances (regions) sorted in descending order and grouped by feature class, illustrating the relative contribution of each temporal–spectral marker to the LSD vs. PLA classification.

Consistent with the global spectra (Figure 2A), LSD induced systematic shifts in oscillatory peak frequencies at the regional level. With the aim of providing clarity on interpretation and on visualization, the first two rows of Figure 3A display the grand-averaged peak shift for alpha and beta bands. This is defined as the difference between the frequency of the LSD peak and the PLA peak in the given band. We showed only the regions that were significant (*p <* 0.05, permutation-corrected) when we compared the LSD peaks to the PLA ones. On average, alpha peaks were shifted upward by approximately 2 Hz and beta peaks by approximately 5 Hz (permutation-corrected, *p <* 0.05). Random Forest classifiers trained on peak-frequency features achieved high decoding accuracy (accuracy *>*82%; Supp. figure S9 for AUC-ROC plots), and the corresponding importance maps closely resembled the spatial patterns observed in the statistical contrasts.

LSD also produced pronounced changes in the aperiodic component of the spectrum. Both the 1/*f* exponent (slope) and offset showed significant, brain-wide effects (Figure 3A, third and fourth rows), characterized by slope flattening and reduced offset. The strongest effects were localized to posterior regions, including the posterior cingulate cortex and visual cortices, as identified by the highest decile of effect magnitudes (*p <* 10*^−^*^3^, magenta). The ML decoding analysis revealed that feature importance generally matched with the regions with the strongest drug effect.

Temporal complexity measures revealed a complementary pattern. Both LZC and HFD increased across the cortex under LSD (Figure 3A, final two rows), with maximal effects again observed in posterior regions (*p <* 10*^−^*^5^, yellow). Decoding models trained on LZC and HFD achieved accuracies of approximately 76% (Supp. figure S9 for AUC-ROC plots), and their feature-importance maps overlapped with regions showing the strongest statistical effects.

Finally, panel 3C extends these analyses using a multidomain Random Forest model that jointly incorporated all temporal and spectral features across cortical regions. Permutation-corrected feature importances revealed that complexity metrics, aperiodic parameters, and spectral power features contributed most strongly to decoding LSD versus PLA. This multidomain ML analysis confirms that LSD-related neural signatures are distributed across multiple feature domains, complementing the single-domain statistical results in Figures 3A and 3B.

### 3.3 Functional decoding reveals differentiated effects across cognitive and affective domains

To relate LSD-induced electrophysiological changes to cognitive function, we performed functional decoding of each neural feature using NiMARE meta-analytic topic maps (Figure 4). Unthresholded LSD–PLA statistical maps were binned into deciles, and each decile was compared against NiMARE topic activation maps to identify which cognitive and affective systems are most affected. Each row of the heatmaps corresponds to a feature family: oscillatory Peak Power (Panels **A, B**), peak-frequency shifts (**C, D**), aperiodic parameters (**E, F**), and temporal complexity measures (**G, H**). The ordering of topic terms reflects their weighted average across deciles (see Section 2.7), and axis directionality is indicated in the figure.

**Figure 4:**
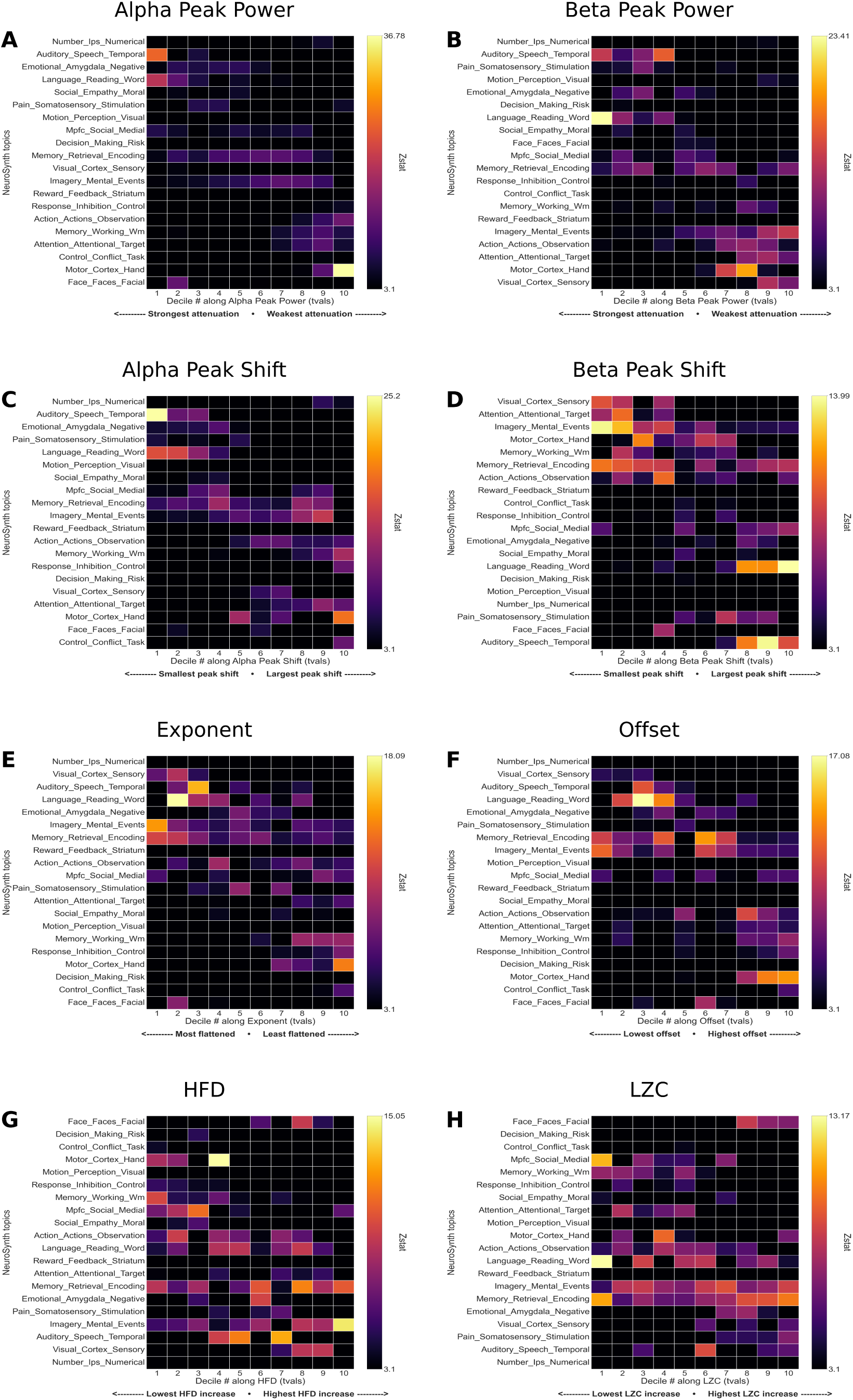
Functional Decoding of LSD-induced electrophysiological changes using NiMARE. Binned (deciles) contrast maps (LSD-PLA) were used to identify how cognitive and affective systems are related to the drug-induced changes. **A** Alpha Peak Power, **B** Beta Peak Power, **C** Alpha Peak Shift, **D** Beta Peak shift, **E** 1/*f* slope, **F** Offset, **G** Fractality (HFD), **H** Complexity (LZC).

The strongest attenuation of alpha Peak Power occurred in auditory, language, and emotion-related systems. Motor and attention regions showed the weakest attenuation, with memory retrieval and imagery networks exhibiting intermediate effects. Domains showing the largest alpha power reductions also exhibited the minimal alpha peak-frequency shifts, while domains with relatively preserved alpha Peak Power (motor, attention) showed larger peak shifts. Consistent with this dissociation, no significant relationship was observed between the magnitude of alpha peak-power reduction and peak-frequency shift (Spearman’s *ρ*, *p >* 0.05), indicating that these effects follow distinct functional hierarchies rather than reflecting a simple inverse relationship.

Beta Peak-Power attenuation displayed a comparable organization, with the strongest reductions in auditory, emotional, and language domains and weaker effects in visual, attentional, motor, and imagery systems. In contrast to the alpha band, beta peak-frequency shifts were largest in domains showing strong beta power attenuation (e.g., auditory and language systems), whereas domains with relatively preserved beta power exhibited smaller frequency shifts.

Aperiodic parameters exhibited a complementary system-level organization. Flattening of 1/*f* slope and the corresponding changes in offset were most pronounced in primary sensory areas (visual and auditory), language-related domains, and higher-order imagery networks, while motor systems showed comparatively minimal effects. Changes in temporal complexity mirrored this pattern: increases in HFD and LZC were largest in mental imagery, visual, auditory, and memory-related domains, and smallest in motor systems. Across domains, systems showing the strongest aperiodic reconfiguration also exhibited the largest increases in temporal complexity.

### 3.4 Exploratory analysis of LSD × Music interaction

We investigated the interaction between LSD and music using a 2×2 factorial design. While no regions survived whole-brain FDR correction, we observed nominally significant interaction effects (p<0.05, uncorrected). These trends point to a negative interaction, where the presence of music appears to attenuate the typical neural signature of LSD (Fig. 5). Specifically, we observed a steepening of the aperiodic slope and increased broadband power—effects that run counter to the spectral flattening typically induced by psychedelics. Similarly, signal complexity metrics (LZC and HFD) indicated that the combination of LSD and music resulted in reduced complexity and fractality relative to LSD in silence, suggesting that auditory stimuli may constrain the effects of the drug.

**Figure 5:**
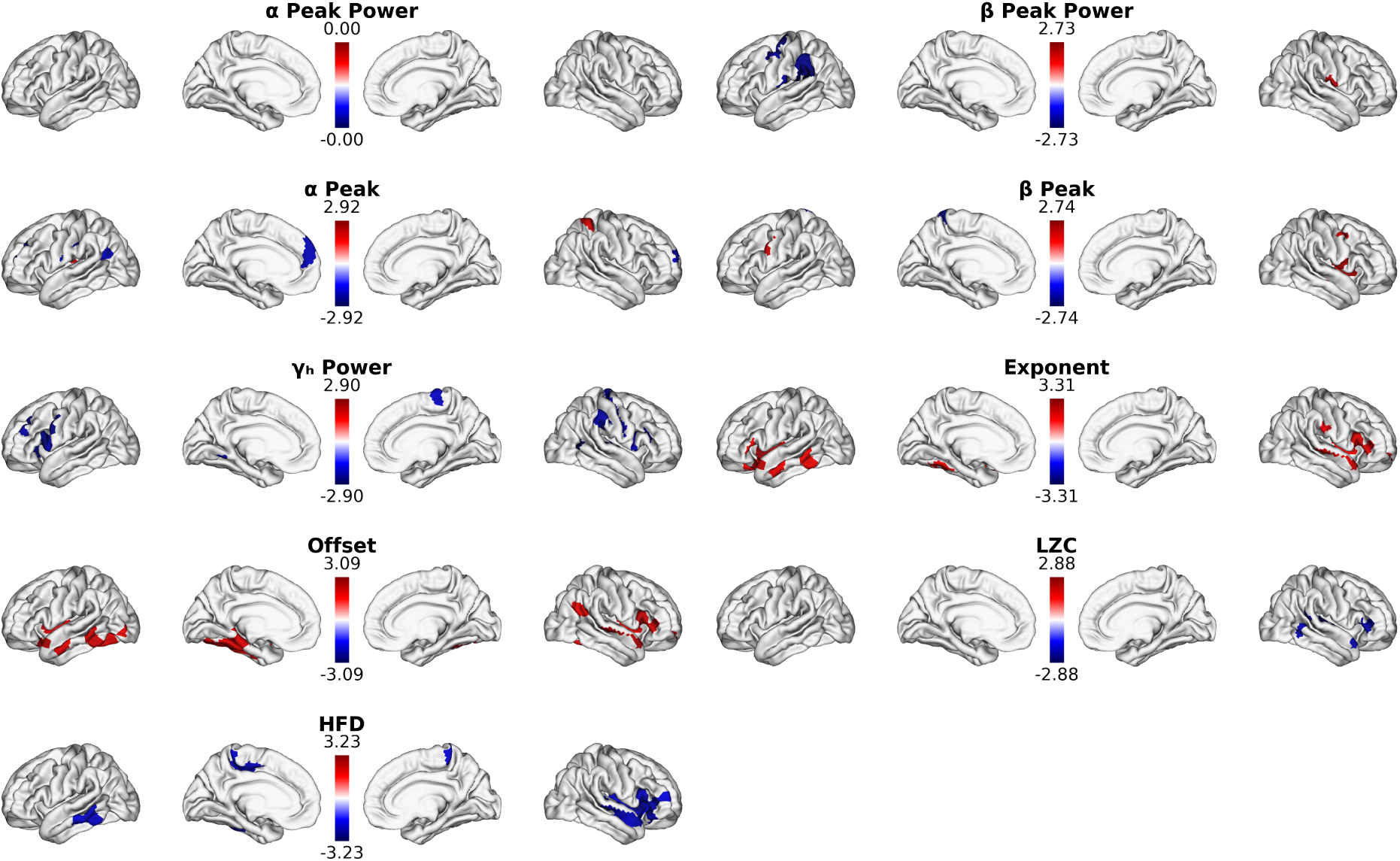
Interaction effects of drug x music. Statistical contrasts are presented for the different spectral and temporal features (*p <* 0.05, uncorrected).

## 4 Discussion

Psychedelics inducing desynchronization in alpha and beta is a hallmark effect that numerous studies have reported about [14, 12, 13]. A limited number of studies have decomposed the power spectrum in its periodic and aperiodic components to show that psychedelics do indeed weaken the oscillatory activity [22, 13]. Evidence for LSD-related increases in alpha and beta frequencies has been noted in the PSDs across few studies [12, 22], but these effects have not yet been systematically quantified or spatially characterized. If LSD speeds up the alpha and beta activity, interpreting the alpha and beta oscillatory weakening risks misinterpretation, as systematic shifts without significant weakening of power within canonical frequency range will masquerade as desynchronization. By focusing on the pure oscillatory mode, we showed robust shifts alpha and beta across brain regions. By analyzing changes in alpha and beta power defined using oscillatory peaks while factoring the Peak shifts into account, we have evidence to suggest that LSD induces genuine weakening of alpha and beta power. This study overall shows LSD induces both weakening of power and at the same time increases alpha and beta peak frequencies. In addition, we investigated the changes in spectral and temporal features as a result of the interaction between LSD and music, building on the previous studies [41]. Our analyses did not identify a statistically significant interaction after correction; however, nominal trends in the exponent, offset, HFD, and LZC point to a relative attenuation of these neural effects during music listening.

### 4.1 LSD elicits localized reductions in alpha and beta oscillation amplitudes

Current neuropharmacological consensus holds that the hallucinogenic effects of psychedelics are primarily mediated by serotonergic signaling (see [9, 67] for review), acting predominantly – though not exclusively – through 5-HT2*A* receptors [8]. This view is supported by studies showing that pretreatment with the 5-HT2*A* antagonist Ketanserin blocks the subjective and neural effects of these drugs [68]. Although the effects of psychedelics on neural circuitry are still being investigated, converging evidence suggests that the 5-HT2*A* agonism induces an asynchronous mode of glutamate release [69, 70]. This mechanism appears to disrupt the temporal synchrony of neural populations, leading to widespread dysregulation of network dynamics. This receptor-mediated disruption is consistent with the regionally selective broadband desynchronization observed in our data, particularly in cortical regions with high 5-HT2*A* receptor density. Such broadband desynchronization is among the most consistently reported electrophysiological signatures of various psychedelics [14, 71].

Across serotonergic psychedelics, attenuation of alpha and beta oscillatory power is a consistent finding, reported for N,N-DMT [13, 72, 17], 5-MeO-DMT [73], and psilocybin [14]. In N,N-DMT studies, increases in low-frequency activity (delta/theta) are not consistently observed and depend on analytical approaches such as the separation of oscillatory and aperiodic components [13]. Consistent with this literature, LSD induces genuine attenuation of oscillatory power. However, we additionally observe robust increases in alpha and beta peak frequencies, indicating a spectral acceleration rather than a replacement of faster rhythms by slower ones. These effects are partly dissociable in space, suggesting distinct but co-occurring mechanisms of cortical reorganization. By comparison, ketamine shows a different electrophysiological profile, with increased gamma activity and reduced lower-frequency coordination [74]. Overall, while reduced oscillatory power may be a shared feature across altered states, LSD is distinguished by a systematic upward shift in alpha and beta peak frequencies. Cross-drug comparisons should nevertheless be interpreted cautiously given differences in modality and analytical methods.

Importantly, electrophysiological recordings reflect a mixture of rhythmic (synchronized) and arrhythmic background activity rather than purely oscillatory processes [75]. The latter typically follows a scale-free, 1/*f* -like distribution [76]. Analyzing raw spectra without separating periodic and aperiodic components risks misinterpretation, because apparent power reductions can arise either from genuine oscillatory desynchronization or from changes in the arrhythmic background that lower total power without altering synchrony. Therefore, distinguishing true oscillatory desynchronization requires explicit parameterization of the spectrum [20]. Recent analytical frameworks have been developed to parameterize and disentangle oscillatory peaks from the aperiodic background [77, 20]. Studies adopting this approach reported an LSD-induced weakening of oscillatory activity [22, 13].

Applying this approach, we show that LSD-induced oscillatory weakening is neither uniform across frequencies nor spatially homogeneous. At the global level (Fig. 2A), spectra replicate the well-established pattern of low-frequency desynchronization both before and after correction for the aperiodic component, attesting to the robustness of our preprocessing pipeline. In raw (non-parametrized) spectra at the regional level, however, relatively few regions exhibit significant LSD–PLA differences. Spectral parameterization reveals that the true oscillatory components are present primarily in alpha and beta and additionally in high-gamma (Supplementary Fig. S1).

Accounting for LSD-induced shifts in oscillatory peak frequencies is critical for interpreting changes in power. Although peak frequency shifts under LSD have been noted previously [12, 22], their implications for power analyses have not been examined in detail. Analyzing power within canonical frequency ranges may mischaracterize oscillatory amplitude when peaks shift; we therefore propose measuring power directly at oscillatory peaks (”Peak Power”). This peak-based analysis (Fig. 2B-C, second row) revealed clear, spatially extended reductions in alpha and beta oscillatory amplitude under LSD. In high-gamma, raw spectra suggested increased power under LSD, whereas parameterized spectra revealed focal power suppression, particularly in posterior regions (Fig. 2D, first and second row). This discrepancy is consistent with a flattening of the aperiodic slope under LSD, which shifts raw spectral power toward higher frequencies without reflecting increased oscillatory activity. A striking illustration of this dissociation was observed in the posterior cingulate cortex (PCC), a core hub of the default mode network with disproportionately high 5-HT2*A* receptor expression [11]. In raw spectra, the PCC exhibited robust LSD–PLA differences across frequency ranges. These differences disappeared after removal of the aperiodic component, indicating that they were driven primarily by changes in the 1/*f* background rather than by oscillatory power per se. Across the cortex, LSD consistently flattened the aperiodic slope, with the strongest effect (*p <* 10^−5^) localized to the PCC. This finding underscores that apparent power reductions in key hub regions may reflect shifts in background spectral properties, rather than true oscillatory desynchronization. In this sense, spectral parametrization does more than merely correct for confounds, but also facilitates interpreting psychedelic effects in physiologically meaningful terms.

One possible interpretation of this flattening is that it reflects changes in the temporal coordination of neural population activity. Prior theoretical, computational, and empirical work suggests that the aperiodic exponent is sensitive to excitation–inhibition balance and population synchrony, with flatter spectra potentially reflecting more irregular, less synchronized population activity [78, 79, 21]. In the context of psychedelics, this interpretation is compatible with evidence that 5-HT2A receptor activation modulates cortical excitability and glutamatergic signaling, including asynchronous glutamate release in prefrontal pyramidal neurons [69, 14]. However, the precise physiological basis of LSD-induced aperiodic changes in human MEG remains to be established.

To further illustrate the consequences of ignoring peak frequency shifts, we also examined band-limited power within canonical frequency ranges after removal of the aperiodic component (Supplementary Figure S6). This conventional approach yielded more spatially extensive apparent desynchronization, particularly in alpha, compared with peak-based analyses. Such differences indicate that fixed-band analyses can inflate the apparent spatial extent of power changes when peak frequencies shift, reinforcing the need for caution when interpreting band-limited power under LSD.

The frequency specificity of LSD-induced power suppression aligns with known neurophysiological principles. Low-frequency oscillations reflect large-scale, spatially distributed synchronization, whereas higher-frequency rhythms tend to arise from more localized circuit interactions [80, 81]. A drug that perturbs global coordination, as serotonergic psychedelics do through 5-HT2*A*-mediated disruption of recurrent processing, would be expected to preferentially weaken large-scale, lower-frequency rhythms. In this context, the topography and frequency specificity of oscillatory suppression observed here are consistent with mechanistic accounts of psychedelic action on cortical hierarchy. Given the close association of these rhythms with top-down signaling [82], their suppression under LSD is consistent with the REBUS (Relaxed Belief under Psychedelics) framework [83].

Complementing these inferential results, machine-learning analyses provided convergent evidence for the robustness of low-frequency oscillatory suppression. Models trained on oscillatory power features achieved high discriminative accuracy between LSD and PLA, with feature importance maps highlighting posterior cortical regions that closely overlapped with those identified by classical statistics. This convergence across independent analytical approaches supports the reliability of the observed effects.

Finally, high-gamma oscillatory activity exhibited a distinct, more focal pattern. Unlike alpha–beta suppression, the effects in the gamma band are not consistently reported in the literature, in part because the gamma activity is highly susceptible to muscle artefacts and therefore often excluded from the analysis. To our knowledge, this study provides the first evidence isolating the oscillatory gamma response to psychedelics. Spectral parametrization (Supplementary Figure S1) confirmed a considerable amount of oscillatory activity in high-gamma. We found that LSD produced a focal decrease in high-gamma, primarily in the posterior regions, including occipital cortex. Previous studies have reported both increase [84] and decrease in gamma activity under psychedelics [14], but those observations were based on raw (non-parametrized) spectra (i.e. with 1/*f* component), which confounds interpretations. In contrast, our results reflect strictly oscillatory power. Notably, classifiers trained exclusively on high-gamma oscillatory power achieved robust decoding performance, with spatially coherent feature importance profiles. If these effects were driven predominantly by electromyographic contamination, one would expect diffuse or inconsistent spatial patterns, which was not observed. While caution remains warranted given the susceptibility of high-frequency activity to artefacts, the convergence of parameterized spectral and decoding results suggests that the observed high-gamma effects likely reflect condition-related neural activity.

### 4.2 LSD increases alpha and beta peak frequencies

Alpha oscillations are among the most extensively studied rhythms in human electrophysiology, dating back to Hans Berger’s earliest EEG recordings [85]. Peak alpha frequency (PAF) has been shown to track attentional state, arousal, and broader cognitive performance [86]. Conceptually, the alpha rhythm has been described as a “fulcrum” of global brain state transitions, reflecting the natural resonant frequency of the resting brain and the stability of cortical-thalamic loops [87]. Because alpha oscillations play a central role in sensory processing, thalamocortical communication, and the regulation of inhibition and excitation balance, even modest shifts in their peak frequency carry mechanistic significance, as they imply changes in the basic operating frequency of cortical networks.

By isolating oscillatory components via spectral parameterization, we show that LSD induces consistent upward shifts in alpha and beta peak frequencies across widespread cortical regions (Figures 2 and 3). This effect differs from those of anaesthesia, which reliably slow alpha rhythms, a pattern associated with reduced cortical excitability and strengthened top-down constraints [88]. Psychedelics, by contrast, generally relax high-level constraints while increasing cortical excitability, and the alpha and beta peak frequency speeding observed here is consistent with this mechanistic dissociation. Notably, comparable peak-frequency modulation has not been robustly observed for psilocybin or DMT, which predominantly flatten oscillatory power without clear peak modulation [89, 13]. Thus, under LSD, reductions in alpha and beta power co-occur with a shift of these rhythms toward faster operating frequencies.

What neuropharmacological features might account for this effect? Several features distinguish LSD from other serotonergic psychedelics. First, LSD exhibits markedly higher affinity for 5-HT2*A* receptors than psilocybin or DMT [8]. Second, crystallography and pharmacology studies reveal that LSD “shuts the lid” on 5-HT2A/2B receptors, giving an exceptionally long residence time, more than twice that of psilocybin [90, 91]. Furthermore, LSD induces stronger increases in thalamocortical connectivity compared with other serotonergic drugs [92]. Given that the alpha-band dynamics depend on factors including cortico-thalamo-cortical connectivity and intracortical circuits mechanisms [93], one plausible, though necessarily speculative, interpretation is that LSD’s unique pharmacokinetic and circuit-level effects simultaneously (a) disinhibit alpha and beta oscillatory generators, (b) alter local excitation/inhibition balance in ways that retune these rhythms to faster frequencies, and (c) amplify thalamocortical drive, shifting the natural resonant frequency of the system. Although robust peak-frequency shifts were observed only under LSD, cross-psychedelic studies with larger cohorts, as well as studies elucidating neuropharmacological similarities and distinctions are both necessary to better establish the specificity of this effect.

Thalamocortical connections serve as a central relaying architecture linking sensory input with cortical hierarchies. LSD-induced increases in thalamocortical connectivity have been interpreted as reduced sensory gating and enhanced bottom-up flow, consistent with alterations in functional hierarchy [92]. Within the REBUS framework, which integrates the entropic brain hypothesis [53] and the free-energy principle [94], psychedelics transiently relax high-level (top-down) priors, weakening top-down predictions while facilitating bottom-up information flow (i.e., less sensory gating). In this context, the combined weakening and speeding-up of alpha and beta rhythms may reflect a hierarchical rebalance: weakening of power indicates a loosening of top-down constraints, while peak-frequency acceleration might indicate heightened thalamic drive (bottom-up flow) and increased cortical excitability. Together, these effects align with the hierarchical reconfiguration proposed by the REBUS framework [83].

### 4.3 LSD exerts system-selective effects

Although LSD induces both oscillatory power reductions and peak-frequency increases across the cortex, the relative expression of these effects differs markedly across functional systems. To characterize this system specificity, we assessed the functional impact profiles of each effect using meta-analytic decoding based on NiMARE. This approach leverages large-scale neuroimaging databases to map regional neurophysiological changes onto functional domains, allowing effect magnitudes to be quantified across cognitive systems. Using this framework, we show that the two principal oscillatory effects of LSD – attenuation and acceleration of alpha–beta rhythms – follow distinct functional hierarchies across the cortex.

LSD-induced suppression of alpha and beta power was strongest in sensory, language, and emotion-related systems, with comparatively minimal effects in motor networks. In particular, auditory, visual, and language systems exhibited pronounced alpha suppression. This pattern is consistent with the high expression of 5-HT2*A* receptors in sensory and multimodal association cortices [11] and aligns with prior electrophysiological work showing widespread psychedelic-induced desynchronization, especially within default mode network hubs such as the posterior cingulate cortex [14]. Emotion-related networks similarly showed strong alpha–beta attenuation, consistent with evidence that LSD modulates limbic and affective circuits and alters emotional processing [95, 96]. In contrast, motor systems remained largely preserved, exhibiting minimal oscillatory suppression. This pattern corroborates the recent connectivity findings that LSD “flattens” the brain’s functional hierarchy by reducing the differentiation between transmodal (e.g. default mode, frontoparietal) and unimodal (primary sensory) cortices [97]. Our results provide electrophysiological support for this account: the largest power weakening in transmodal networks (language/emotional hubs) and in unimodal regions (primary sensory areas) reflect a substantial reconfiguration of the normal hierarchical order of oscillatory activity under LSD.

In contrast to power suppression, oscillatory peak-frequency modulation followed a distinct system-level organization. Regions exhibiting the strongest alpha power attenuation—such as auditory and language networks—showed minimal shifts in alpha peak frequency, whereas areas with more preserved alpha power, including motor and attentional systems, exhibited larger frequency increases. In the beta band, this relationship was reversed: systems showing the strongest beta power suppression also displayed the largest beta peak-frequency shifts. These dissociations could indicate two neurophysiological processes –dampening of oscillatory power in some circuits and acceleration of rhythmic activity in others – organized along distinct functional hierarchies. However, our results comparing the association between dampening of oscillatory power and peak frequency shifts revealed no statistical association (region-averaged Spearman’s *ρ* = −0.04 and −0.12 for alpha and beta respectively, p > 0.05 uncorrected). These results show no evidence of a strong association between the two LSD-induced effects, and thus do not support a common spatial covariation. This interpretation is consistent with prior work: Carhart-Harris et. al [12] noted that LSD not only reduced alpha power but was also associated with a slight increase in the dominant alpha rhythm frequency. Although direct literature on frequency shifts remains limited, our observations reinforce the view that LSD modulates not just the power but also the timing of neural oscillations. Further studies are needed to disentangle these co-occurring effects to shed light on their underlying neuropharmacological mechanisms.

Beyond oscillatory components, LSD also produced pronounced, system-selective changes in the aperiodic background of the MEG spectrum. We observed robust flattening of the 1/*f* slope, most prominently in visual, auditory, language, and imagery-related systems. A flatter 1/*f* slope reflects reduced dominance of low-frequency power and a shift toward more broadband, irregular dynamics [79]. In sensory cortices, where 5-HT2*A* receptor density is high, such flattening likely reflects disrupted inhibitory constraints and increased excitability. Notably, higher-order language and imagery networks exhibited comparable slope flattening, whereas motor regions showed minimal changes, reinforcing the graded and system-selective nature of LSD’s effects (Figure 4).

These aperiodic changes were closely mirrored by increases in neural signal complexity, as indexed by Higuchi fractal dimension and Lempel–Ziv complexity. Regions with the strongest 1/*f* flattening (visual, imagery areas, auditory and language cortices) showed the largest increases in neural diversity, whereas motor regions remained relatively stable. This inverse relationship between spectral slope and signal complexity has been reported previously [98] and suggests a shared mechanism whereby broadband, irregular activity gives rise to richer dynamical repertoires. These findings are consistent with the entropic brain hypothesis [53] and align with prior reports of elevated signal diversity under psychedelics [42, 13, 99]. Given that Lempel–Ziv complexity is sensitive to the regularity of oscillatory activity, such that more rhythmic signals (e.g., alpha) tend to yield lower complexity [100], we performed complementary band-specific analyses. These revealed increased complexity in beta and low-gamma bands, with no significant changes in alpha, indicating that the observed LZC effects are not driven by alpha-band dynamics. Rather, they reflect a frequency-specific increase in neural signal diversity, consistent with a reorganization of cortical dynamics beyond simple power changes.

Together, LSD reorganizes cortical dynamics through convergent mechanisms that are differentially expressed across cortical functional hierarchies. Alpha–beta power suppression, peak-frequency modulation, aperiodic flattening, and increased signal complexity co-localize preferentially within sensory, language, emotional, and imagery systems, while motor networks remain comparatively preserved. This system-selective reorganization provides electrophysiological support for hierarchical and entropic models of psychedelic action, including REBUS, by showing that LSD relaxes structured top-down constraints while expanding the dynamical repertoire of perceptual and cognitive circuits.

### 4.4 Why might Music not be interacting with LSD to potentiate effects

One of our central questions was whether music potentiates the neurophysiological effects of LSD. Across all spectral and temporal features examined, the interaction between music and LSD did not survive correction for multiple comparisons and therefore only be considered as a nominally significant effect. This absence of a robust interaction is noteworthy, given that music is widely regarded as a potent emotional and physiological stimulus [25], has been intertwined with psychedelic ritual practices for millennia [7], and is increasingly incorporated into psychedelic-assisted therapy to enhance emotional depth and meaning-making [30, 101].

Although the interaction effects did not survive correction for multiple comparisons, the resulting maps showed a qualitatively consistent pattern indicative of attenuation rather than potentiation. Specifically, LSD x music interaction was associated with increased exponent and offset, alongside reduced LZC and HFD, indicative of steeper aperiodic slopes, higher broadband power, and reduced temporal complexity relative to the NoMusic condition. Prior work provides a useful framework for interpreting this pattern. Mediano and Rosas [41], with the same dataset, showed that psychedelic-induced increases in neural signal diversity are maximal during internally oriented states (e.g. eyes-closed) and diminish when attention is directed toward external stimuli. Our results extend this finding by suggesting that music may act as a form of grounding agent, an external attentional anchor that stabilizes neural activity and counteracts the unconstrained dynamics characteristic of the psychedelic state. Recent work analyzing various tasks during psilocybin sessions demonstrates similar reductions in neural and subjective intensity when attention is externally directed [4]. Together, these findings raise the possibility that music does not amplify the core neural effects of LSD, but may instead attenuate them by imposing structure and redirecting attention outward. Nevertheless, given the absence of interaction effects surviving correction and the fact that the experimental design was not optimized to isolate subtle state–context interactions, these interpretations should be viewed as tentative. Moreover, detecting interaction effects typically requires substantially greater statistical power than main effects [102], and the present study was not specifically powered for this purpose. Accordingly, this finding should be considered exploratory and in need of replication in larger samples.

In exploratory analyses linking LSD-induced neural changes to subjective experience (Supplementary Figure S2), region-averaged reductions in alpha and beta Peak Power were associated with increased Simple Imagery, whereas flatter spectral slopes, lower offset, reduced beta Peak Power, and lower LZC predicted stronger Complex Imagery. In contrast, LSD-induced neural changes did not significantly predict Intensity, Emotional Arousal, or Positive Mood. Ego Dissolution was selectively associated with beta peak-frequency shifts and LZC. Notably, only Emotional Arousal showed a significant association with the LSD × music interaction: smaller beta peak shifts during music listening were linked to higher emotional arousal. Although these associations are exploratory (*p <* 0.05, uncorrected), they converge with prior reports [41] suggesting that engagement with structured external stimuli can dampen core psychedelic neural signatures even when subjective experience remains emotionally salient.

## 5 Limitations and Perspectives

Several limitations of the present work warrant considerations and can help guide future research. First, because we frequently observed multiple oscillatory peaks (Center Frequencies) within alpha band, we defined the peak as the frequency with maximal amplitude in the 1/f-removed spectrum. Our analysis looking at the Center Frequency, we found that alpha and beta peaks were present in nearly all PSDs (Subjects x Regions), and high-gamma peaks appeared frequently as well, with a slight reduction under LSD. Given that oscillations in alpha and beta rhythms were present in most PSDs, our manual peak-identification procedure produces results largely equivalent to the center-frequency approach. Nevertheless, caution is warranted when interpreting findings in the high-gamma range, where oscillatory structure is less consistent and peak estimation is inherently more variable than in the lower frequency range. In addition, the absence of concurrent EMG recordings did not allow for direct statistical comparison of muscular activity between LSD and placebo conditions, highlighting a limitation that future data collection during LSD should address. Second limitation is in regards to nominal significant interaction between LSD and music. While this aligns with prior reports suggesting that the acute influence of psychedelics may wane when participants engage with external stimuli, it also highlights a broader limitation in how such effects are typically modeled. Most studies, including ours, implicitly assume that LSD x music interactions remain constant over time. However, music is a dynamic and richly structured stimulus, and any modulatory effects may be transient or state-dependent. Collapsing the temporal dimension may therefore obscure meaningful fluctuations. A more time-resolved approach may offer clearer insights by enabling joint tracking of neural responses and concurrent phenomenology. In addition, because the pre-assessment ensuring the preference of genre and the music track was conducted with a separate group, the propensity for this cohort was not established. From a statistical standpoint, interaction effects are inherently more difficult to detect than main effects and typically require substantially larger sample sizes to achieve comparable statistical power [102]. This poses a particular challenge in psychedelic research, where studies are often constrained by modest sample sizes, as in the present dataset (N = 17). As a result, the current analysis may be underpowered to reliably detect subtle state–context interactions, and the observed effects should therefore be interpreted with appropriate caution. A further limitation concerns the characterization of individual differences in subjective experience. At the region level, we observed significant correlations between neurophysiological changes and phenomenology only in the Simple Effects conditions. With regards to Main Effect Drug and Interaction effect drug x music, global (region-averaged) measures did show meaningful associations. We regard these global correlations as exploratory because they were not corrected for multiple comparisons. The lack of region-level associations may be due, in part, to the timing of subjective reporting. Participants provided in-scanner ratings relative to their peak drug experience, whereas MEG acquisition occurred approximately 150 minutes after peak effect. This discrepancy likely introduces noise and reduces the sensitivity of brain–phenomenology correlations.

## 6 Conclusion

In this study, we provide a comprehensive electrophysiological characterization of how LSD reorganizes human cortical dynamics, demonstrating that its acute effects arise from dissociable mechanisms that map systematically onto large-scale functional systems. By combining spectral parametrization, peak-resolved oscillatory analysis, aperiodic modeling, and functional decoding, we identify three principal axes of modulation. First, LSD induces a strong attenuation of alpha and beta power in auditory, visual, language, and emotion-related networks, regions rich in 5-HT2*A* receptors, while sparing motor cortex. Second, LSD produces system-specific increase in alpha and beta peak frequencies that follows a distinct functional hierarchy, independent from that of power suppression. Third, LSD flattens the aperiodic (1/*f*) spectrum and increases temporal complexity (HFD, LZC), with the largest changes in sensory and imagery domains, consistent with a shift toward a desynchronized, high-complexity cortical regime. These results address our initial questions: (Q1) peak-frequency shifts are spatially structured; (Q2) oscillatory power weakening, aperiodic flattening, and complexity increases are anatomically selective; and (Q3) music’s modulatory influence is limited and context dependent, showing no uniform enhancement of LSD’s neural effects and in some cases attenuating them. Overall, LSD induces a structured reorganization of cortical dynamics across temporal and spectral properties, rather than a uniform global perturbation. Future work aimed at disentangling the neuropharmacological mechanisms underlying oscillatory power weakening and peak shifts, and establishing tighter alignment between neural activity and subjective experience, will be essential for clarifying how these LSD-specific effects arise and how they contribute to psychedelic phenomenology.

## 7 Ethics Statement

We utilized the subset of the data collected by Carhart-Harris and colleagues [12]. This original study was approved by the National Research Ethics Service Committee London-West London and was conducted in accordance with the revised declaration of Helsinki (2000), the International Committee on Harmonization Good Clinical Practice guidelines, and the National Health Service Research Governance Framework. Imperial College, London sponsored the research, which was conducted under a Home Office license for research with Schedule 1 drugs.

## 8 Data and Code Availability

Queries regarding data are to be directed to SM and RC-H. Code is available on https://github.com/venkateshness/specparam_LSD

## 9 Authors Contributions

**VS** : Conceptualization, Methodology, Software, Validation, Formal Analysis, Investigation, Data Curation, Writing - Original Draft, Writing - Review & Editing, Visualization. **TN** : Conceptualization (Peak Shifts analysis), Writing - Review & Editing. **AP** : Conceptualization (Preprocessing, Source Localization), Methodology (Preprocessing, Source Localization), Writing - Review & Editing. **JB** : Conceptualization (Statistics), Methodology (Statistics), Writing - Review & Editing. **YJMR** : Conceptualization (Preprocessing), Methodology (Machine Learning), Writing - Review & Editing. **YH** : Conceptualization (Preprocessing), Writing - Review & Editing. **SM** : Data Curation, Writing - Review & Editing. **RC-H** : Data Curation, Writing - Review & Editing. **GL** : Conceptualization, Writing—Review & Editing, Supervision, Project Administration; **NF** : Conceptualization, Writing—Review & Editing, Supervision, Project Administration. **KJ** : Conceptualization, Methodology, Resources, Writing—Review & Editing, Supervision, Project Administration, Funding acquisition.

## 11 Acknowledgments

We thank several colleagues who contributed to the progress of this study. We are grateful to Jordan O’Byrne for assistance with dataset organization; Kenneth Shinozuka for identifying missing data for two subjects; Arianna Caputo for sharing resources and insights on the Higuchi Fractal Dimension; and Yassine El Ouahidi for adapting the functional decoding pipeline from Neurosynth to NiMARE.

K.J. is supported by funding from the Canada Research Chairs (950-232368) program and a Discovery Grant from the Natural Sciences and Engineering Research Council of Canada (2021-03426).

We utilized a subset of the data collected by Carhart-Harris and colleagues [12]. Data collection of this study was supported by Safra Foundation and the Beckley Foundation as part of the Beckley-Imperial research programme, and by supporters of the Walacea.com crowdfunding campaign.

