## Supplementary Material for "LSD Reconfigures Cortical Dynamics Through Faster Brain Rhythms and Increased Fractal Dimension"

May 29, 2026

### 1 Supplementary Material

#### 1.1 Oscillatory peaks present most reliably in alpha and beta bands

To determine whether oscillatory peaks observed across subjects reflected meaningful structure rather than chance detections, we applied a binomial test against a theoretical false-positive rate of 0.05. For each parcel and frequency band, oscillatory peaks identified by the spectral parametrization model were binarized for each subject (peak present or absent). Because each subject has an estimated 5% probability of exhibiting a spurious peak, the number of detected peaks across our 17 subjects can be modeled as a series of independent Bernoulli trials with a null probability of 0.05.

For each parcel, we counted how many subjects showed a peak and then asked: What is the probability of observing at least this many detections purely by chance? This probability

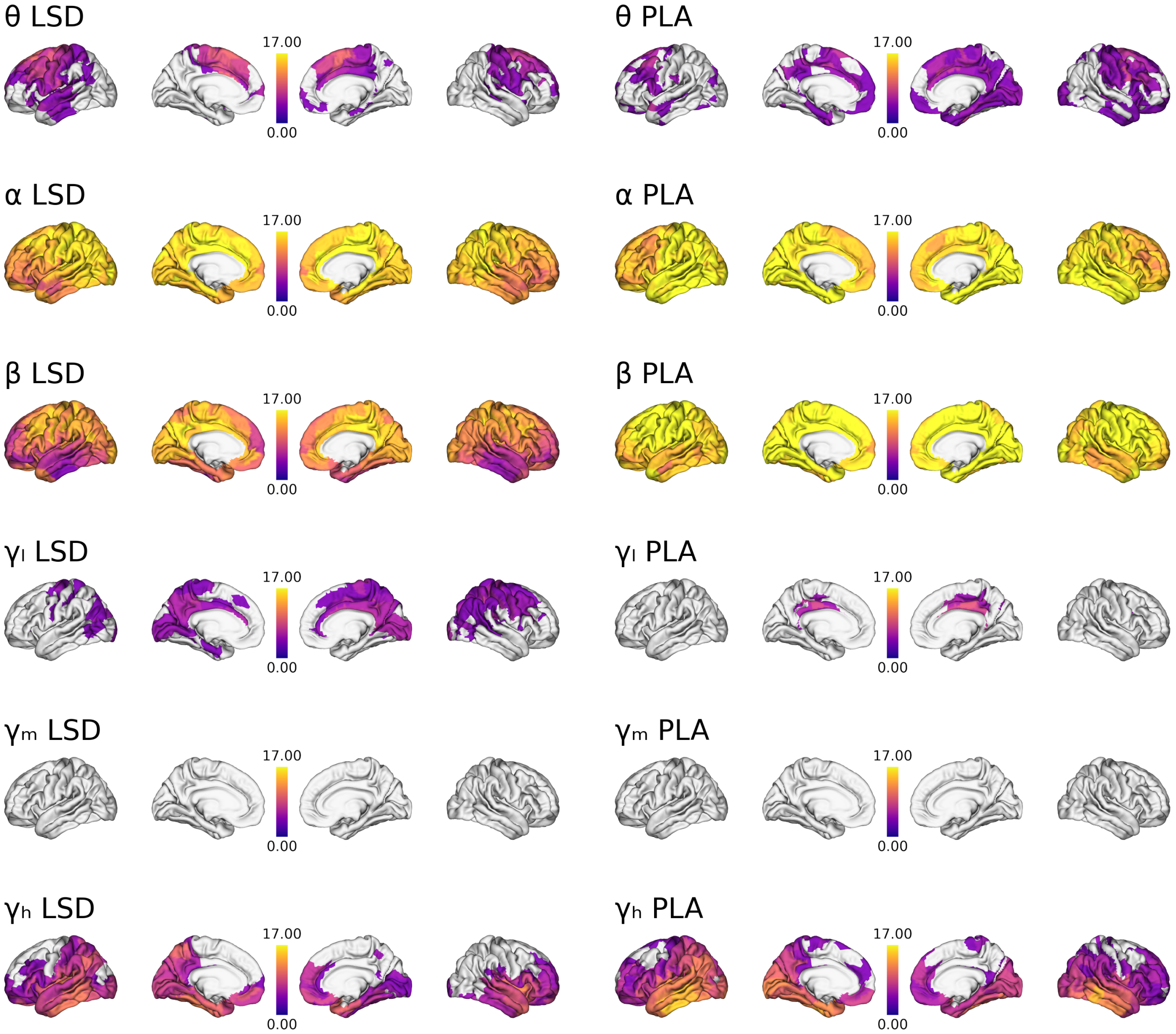


Figure S1: Oscillatory peak counts across frequency bands and drug conditions. For each cortical region, color intensity reflects the number of participants exhibiting a statistically significant oscillatory peak ($p < 0.05$, FDR-corrected) across $\theta$, $\alpha$, $\beta$, low-$\gamma$, mid-$\gamma$, and high-$\gamma$ bands. Left and right halves of each row show peak-count maps for the LSD and placebo (PLA) conditions, respectively. Warmer colors indicate a greater number of participants with detectable peaks in that band.

is given by the upper tail of the binomial distribution (also known as the survival function), which quantifies how extreme the observed count is under the null model of noise-driven peaks. The resulting p-values were then corrected for multiple comparisons across parcels using FDR.

This procedure, adapted from previous work [1], provides a principled way to threshold the Oscillatory Peak Count maps by evaluating whether the across-subject consistency of peak detection exceeds what would be expected from false positives alone (Supplementary Fig S1).

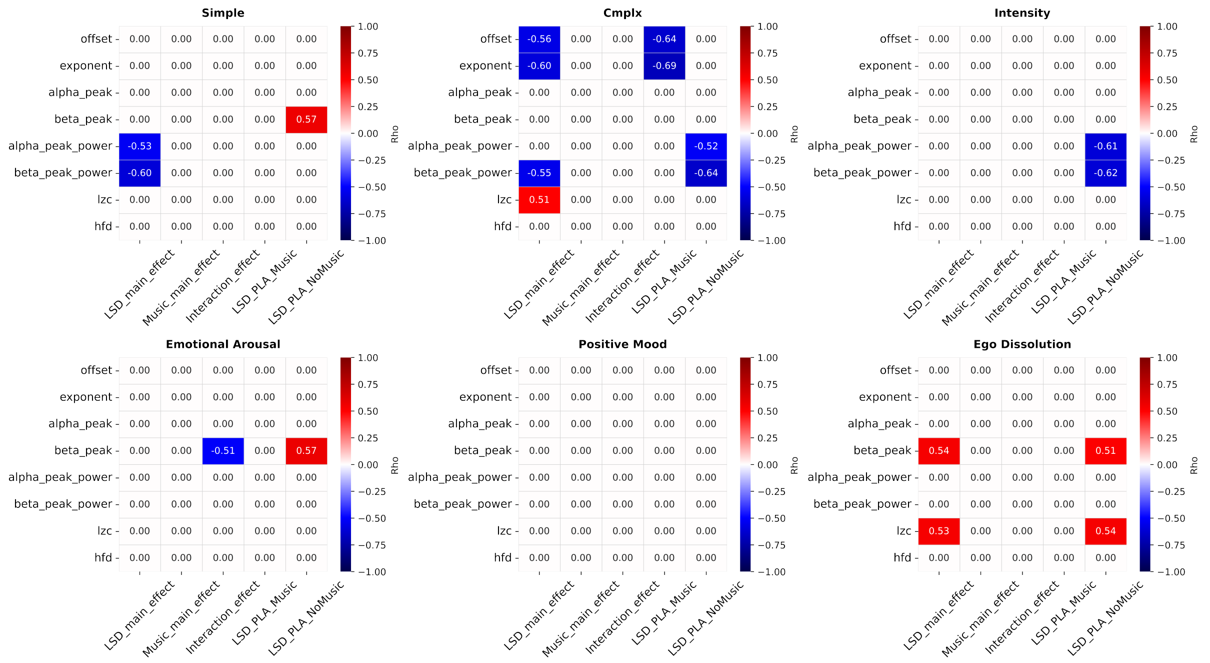


Figure S2: Global-level Regression analysis between electrophysiological features and Phenomenology.

### 1.2 Relations of the temporal and spectral features with Phenomenological dimensions at the global scale

How changes in the neurophysiological features relate to phenomenology is an important question that we addressed by regression analyses. To begin with, we assessed how the changes in the features *global level* i.e., region-averaged, relate to six phenomenological dimensions : 1) Simple Imagery; 2) Complex Imagery; 3) Intensity; 4) Emotional Arousal; 5) Positive mood; 6) Ego Dissolution. These analyses have been conducted for various statistical conditions: Main Effect Drug, Main Effect Music, Interaction Effects, Simple Effect Music, Simple Effect NoMusic. The corresponding heatmaps are presented in the supplementary material (Supplementary Figure S2). The raw phenomenological scores are provided also in the supplementary material (Supplementary Figure S5). Spearman's $\rho$ characterizes the level of correlation between the feature and the subjective ratings, ranging from -1 to 1. We can observe a tight link between feature phenomenology across many phenomenological dimensions. For example, flattening of the slope, and lowering of the offset were significantly associated with increased Complex Imagery. This trend was observed under the Main Effect Drug, and Simple Effect Music contexts. Similarly, alpha and beta Peak Power attenuation significantly correlated with drug's Intensity under Simple Effect NoMusic. In addition, increased LZC was associated with increased Complex Imagery under the Main Effect Drug. This gives insights about the trend at the global level.

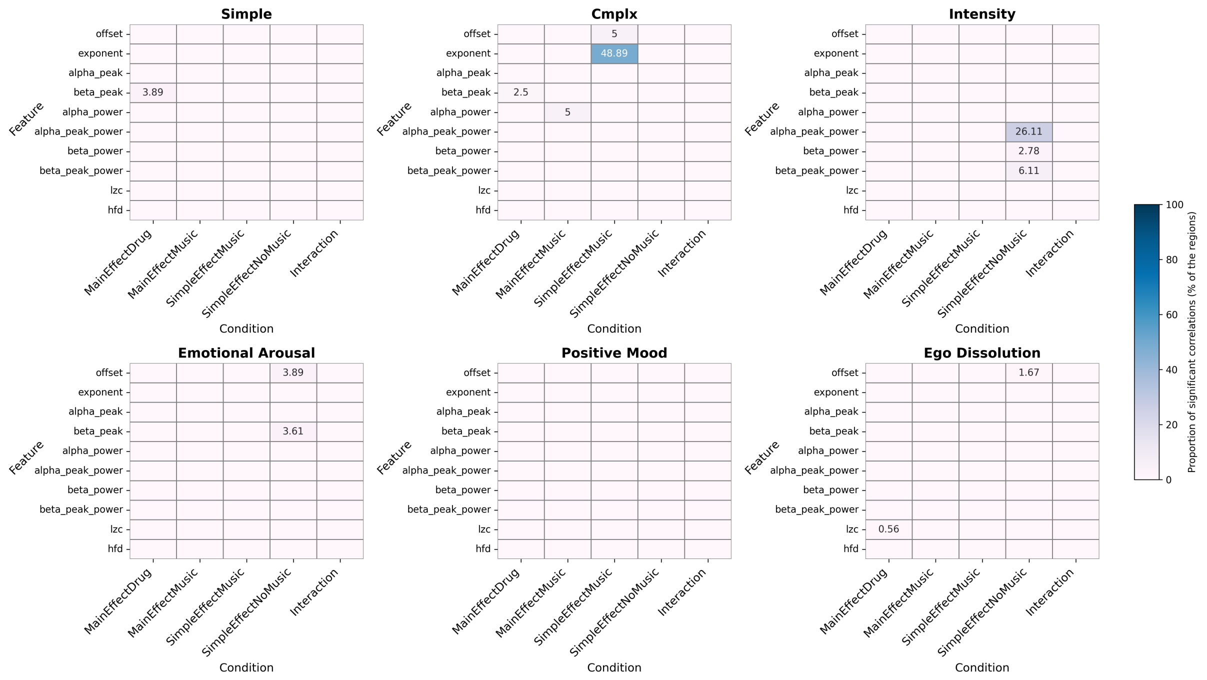


Figure S3: Proportion of Regions with Significant Regressions by Phenomenology Dimension.

### 1.3 Proportion of Regions with Significant Regressions by Phenomenology Dimension

We also investigated the region-level dynamics by performing regression analyses at a finer spatial scale (ROI-level). To this end, we ran Spearman's $\rho$ tests across all the regions (N=360), and the significant ($p < 0.05$; FDR-corrected) correlation results are presented in the supplementary material (Supplementary Figure S3). Proportion of the significant correlation ($p < 0.05$, FDR-corrected) following the Spearman's $\rho$ comparing the changes in neurophysiology at the individual region level and Phenomenology. Units here represent proportions (%) i.e. how many regions are significantly linked to phenomenology out of all regions. Main Effect Drug and Interaction effects are the conditions of interest. Shifts in beta peaks significantly correlate with increased imagery (both simple and complex) in less than 5% of the regions.

### 1.4 Similar topography in spectral features when fitting Specparam up to 120Hz and upto 45Hz

To assess the robustness of our spectral parameterization models, we additionally fit the models using a narrower frequency range (1–45 Hz; Supp. Fig. S4). The first row summarizes model performance on the global spectra, with violin plots showing the distribution of $R^2$, fitting error, offset, and exponent across subjects and conditions. The lower panels display the significant contrast maps for Peak Power and Peak Shifts derived from the narrowband fits. Notably, these maps exhibit highly similar spatial patterns to those obtained with the full 1–120 Hz model, indicating that our main findings are stable with respect to the chosen frequency range and not driven by high-frequency components.

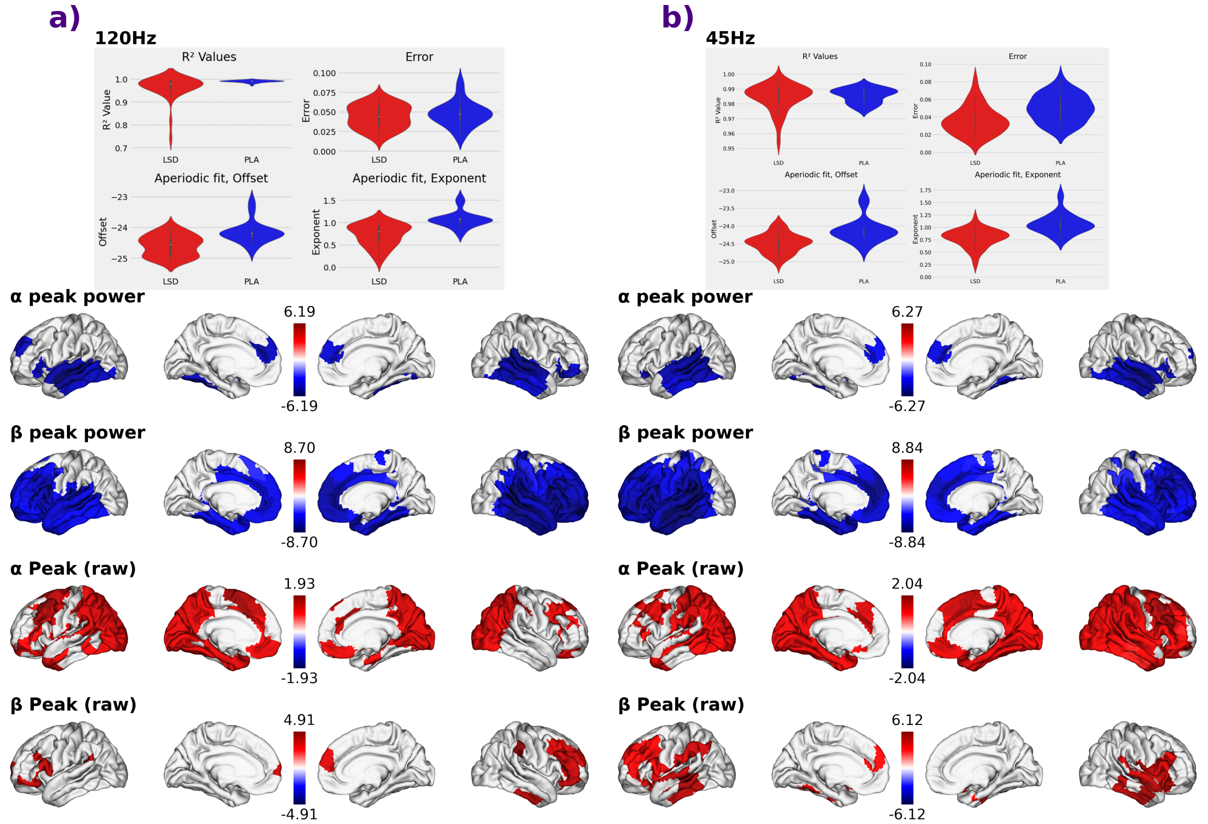


Figure S4: Performance of spectral parameterization models across frequency ranges. **a-b** Global (region-averaged) FOOOF model performance for the broadband (1–120 Hz; a) and narrowband (1–45 Hz; b) spectral ranges. Violin plots show model fit quality ($R^2$ and error) and aperiodic parameters (offset and exponent) under LSD and PLA. Error rate, $R^2$, Offset and Exponent of the global (i.e. region-averaged PSD) specparam model (top row) for the frequency range of 1-120Hz (**a**, same as in the manuscript), and for the narrower frequency of 1-45Hz (**b**). Spatial maps under the barplot show the corresponding LSD-PLA contrast maps for $\alpha$ and $\beta$ Peak Power changes, alpha and $\beta$ peak shifts across two frequency ranges. Maps show the magnitude and direction of LSD–PLA contrasts, illustrating that the qualitative spatial patterns of peak-power suppression and peak-frequency shifts are replicated across both parametrization ranges.

### 1.5 Subjective Reports data

We present here the raw phenomenological scores across the drug and task conditions (Supplementary Fig. S5). The scores were statistically compared using paired t-tests, followed by FDR-correction to account for all the pairwise comparisons.

### 1.6 Effect of analyzing the power dynamics within canonical frequency range

LSD induces both a desynchronization of oscillatory activity and an acceleration of the alpha and beta rhythms. A key methodological contribution of this work is the mitigation of confounds that arise when power is analyzed strictly within canonical frequency bands, without accounting for drug-induced peak shifts. To address this, we quantify oscillatory amplitude using Peak Power, which extracts the power specifically at the individualized peak frequency. This approach isolates genuine oscillatory changes from those driven by shifts in peak location.

Panels (a) and (b) in Supp. fig. S6 show statistically thresholded contrast maps (using the same permutation-based pipeline as in the main text) for alpha and beta Peak Power, respectively. Significant contrast maps (same statistical pipeline as in the main text) for alpha (a) and beta (b) are presented with $1/f$ (first row) and without (second row).

### 1.7 Changes in spatial hierarchy between LSD and PLA

To examine whether the observed effects were spatially global or constrained along cortical hierarchies, we evaluated each region's position within the spatial distribution of effects (Supp. Fig. S7). For every subject and condition, regional values were spatially z-scored to remove global amplitude differences and emphasize relative hierarchical patterns. We then conducted two-tailed two-sample permutation tests to assess the Main Effects Drug and the Interaction (Drug x Music). Contrast maps for the Main Effects were thresholded at $p < 0.05$ and corrected for multiple comparisons using 50,000 permutations, whereas Interaction maps are displayed uncorrected due to their weaker effect sizes.

### 1.8 Band-specific LZC reveals increased complexity in beta and low-gamma

We computed LZC for each frequency band starting with alpha, going up to high-gamma (Supp. Figure. S8).

### 1.9 AUC-ROC reveals strong sensitivity and specificity of the ML models

In addition to reporting the accuracy and feature importances of the ML models, we report here the AUC-ROC (Supp. Figure. S9).

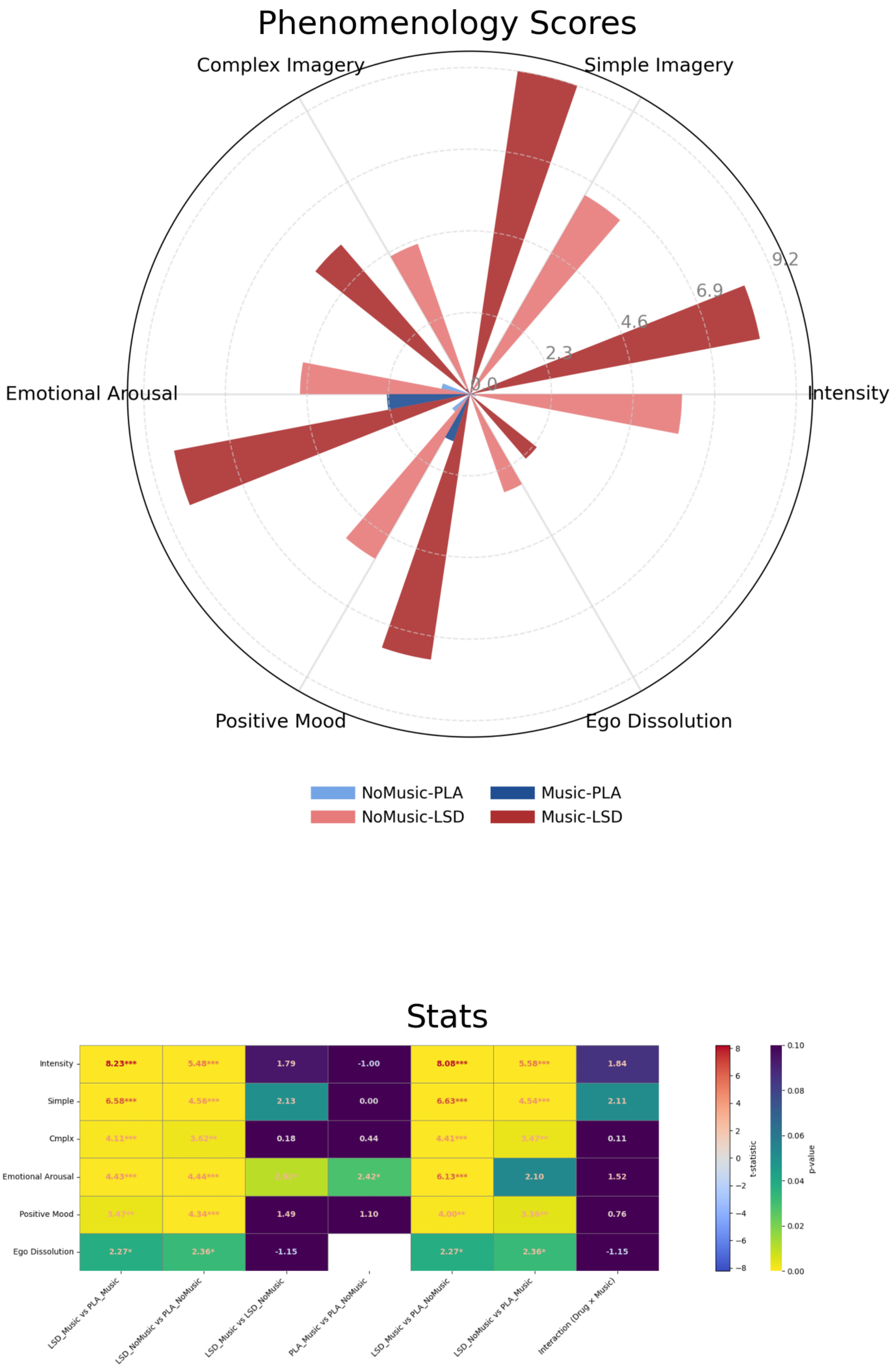


Figure S5: In-scanner subjective ratings across conditions. First and second row respectively correspond to grand-averaged scores and the paired statistical comparisons between LSD and PLA conditions ($p < 0.05$, FDR-corrected).

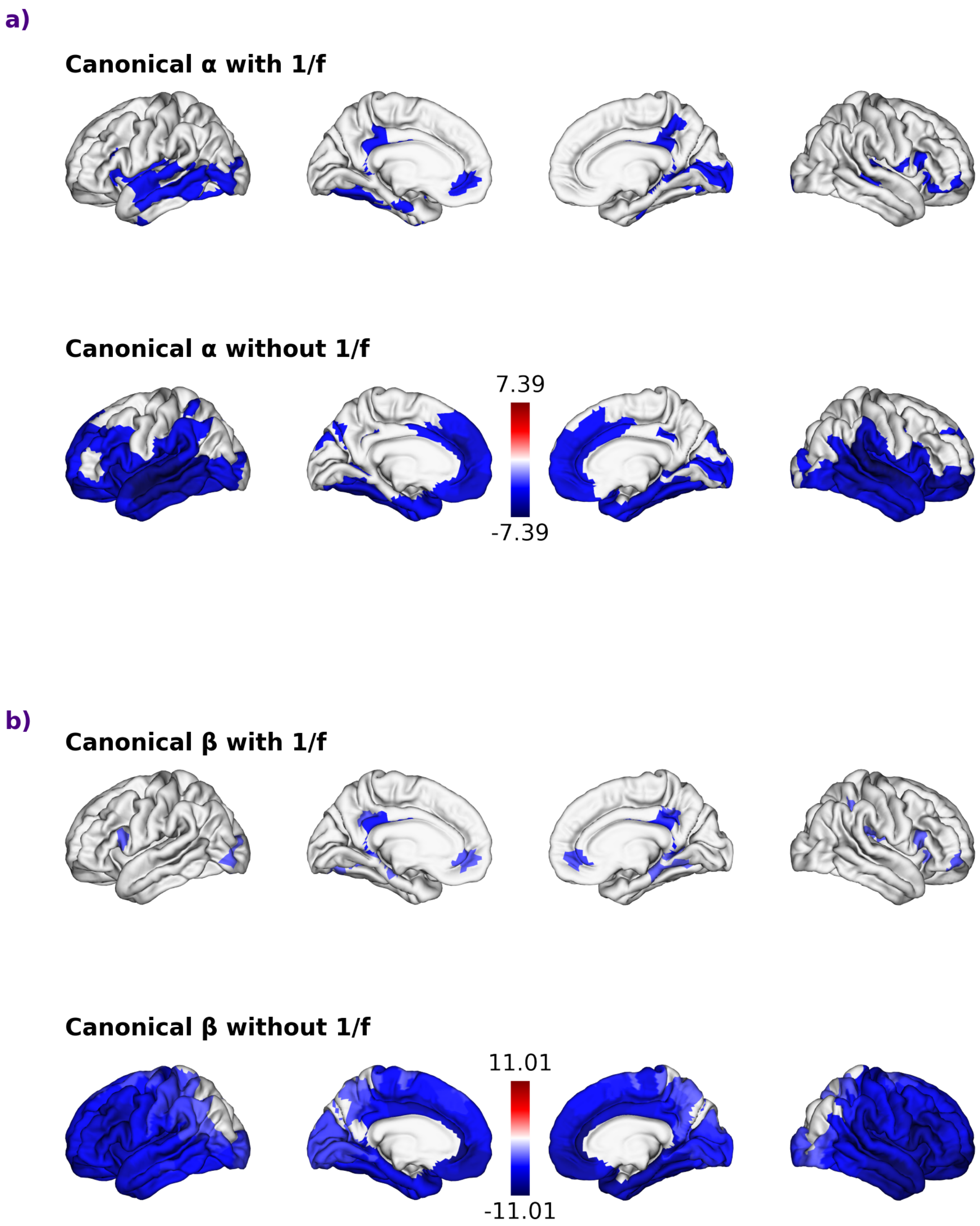


Figure S6: Power changes when analyzing with peak shifts. **a-b** Region-resolved changes in bandlimited power between LSD and PLA ($p<0.05$; permutation-corrected). For each band, the top row shows canonical band-power differences with the aperiodic ($1/f$) component included, and the bottom row shows differences after removing the $1/f$ component (oscillatory-only power).

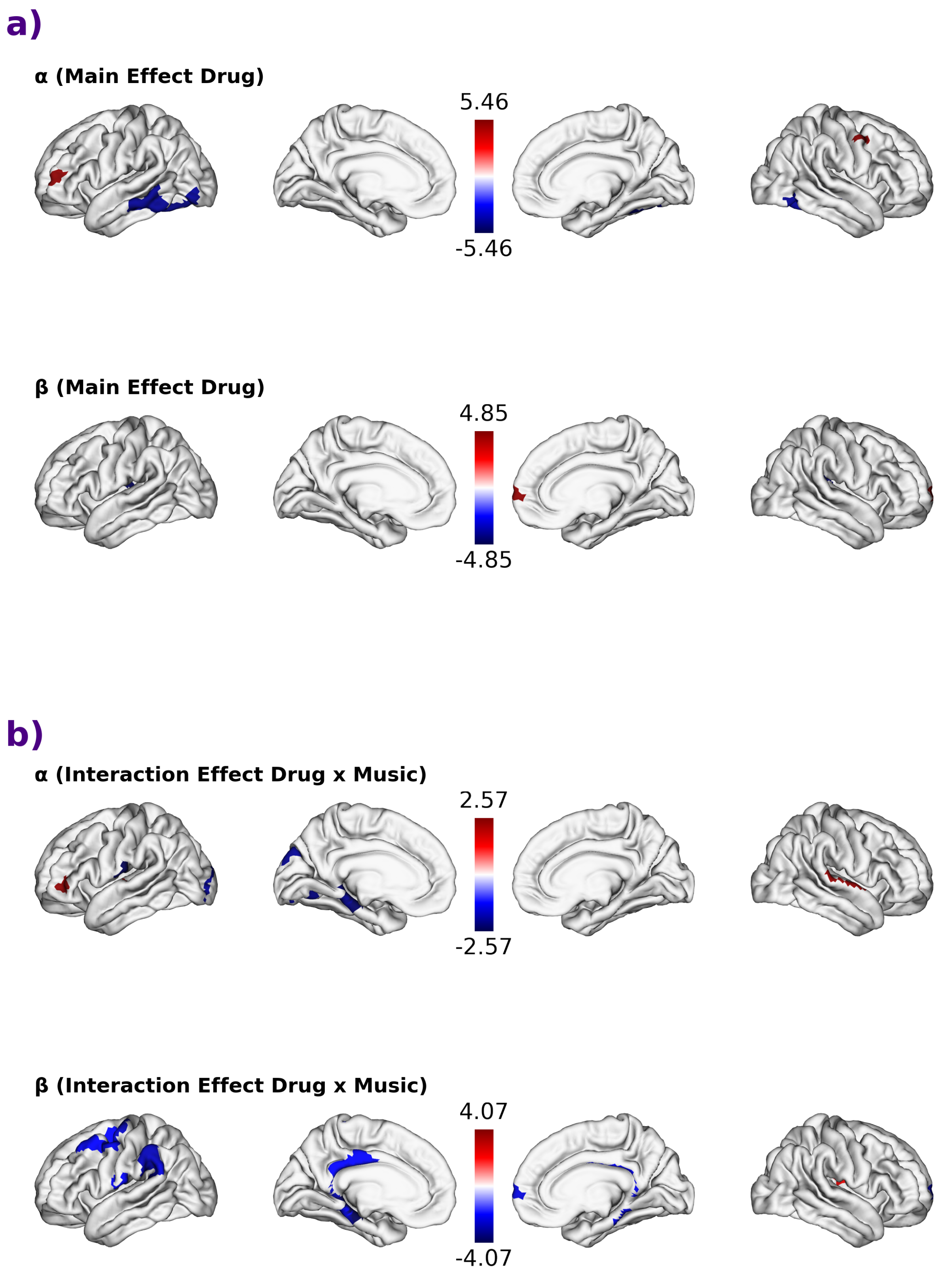


Figure S7: Changes in spatial hierarchy between drug conditions. **a** Spatial z-scored main effects of Drug (LSD vs PLA) for $\alpha$ and $\beta$ Peak Power. For each subject and condition, regional values were spatially z-scored to emphasize hierarchical (relative) patterns rather than global amplitude differences. Shown are permutation-corrected contrast maps (two-tailed; 50,000 permutations; $p < 0.05$). Red or Blue: LSD pushes the given region a spatially important place out of the rest. **b** Spatial z-scored interaction effects of Drug × Music for $\alpha$ and $\beta$ power. Interaction maps are displayed uncorrected due to their weaker and more spatially localized effects. These patterns suggest that music modulates LSD-induced oscillatory suppression primarily in temporal and parietal association cortices. Red or Blue : LSD and Music together push the given region out of the rest

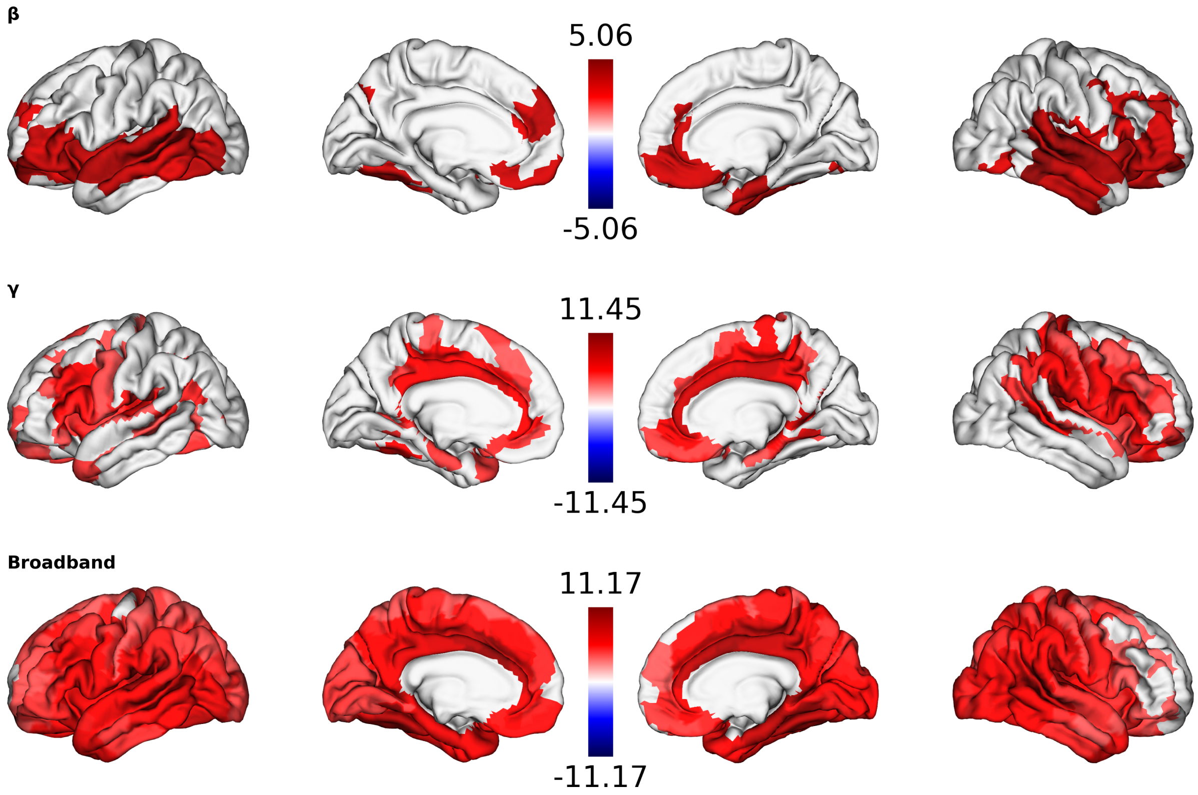


Figure S8: Group-level differences in adaptive Lempel-Ziv Complexity. For context, the significant (permutation-corrected, N=50,000, p<0.05) map for broadband signal (as in manuscript) is on the left. LZC computed on the filtered recordings showing significance in beta and low-gamma activity, while showing no significance in alpha, mid- and high-gamma.

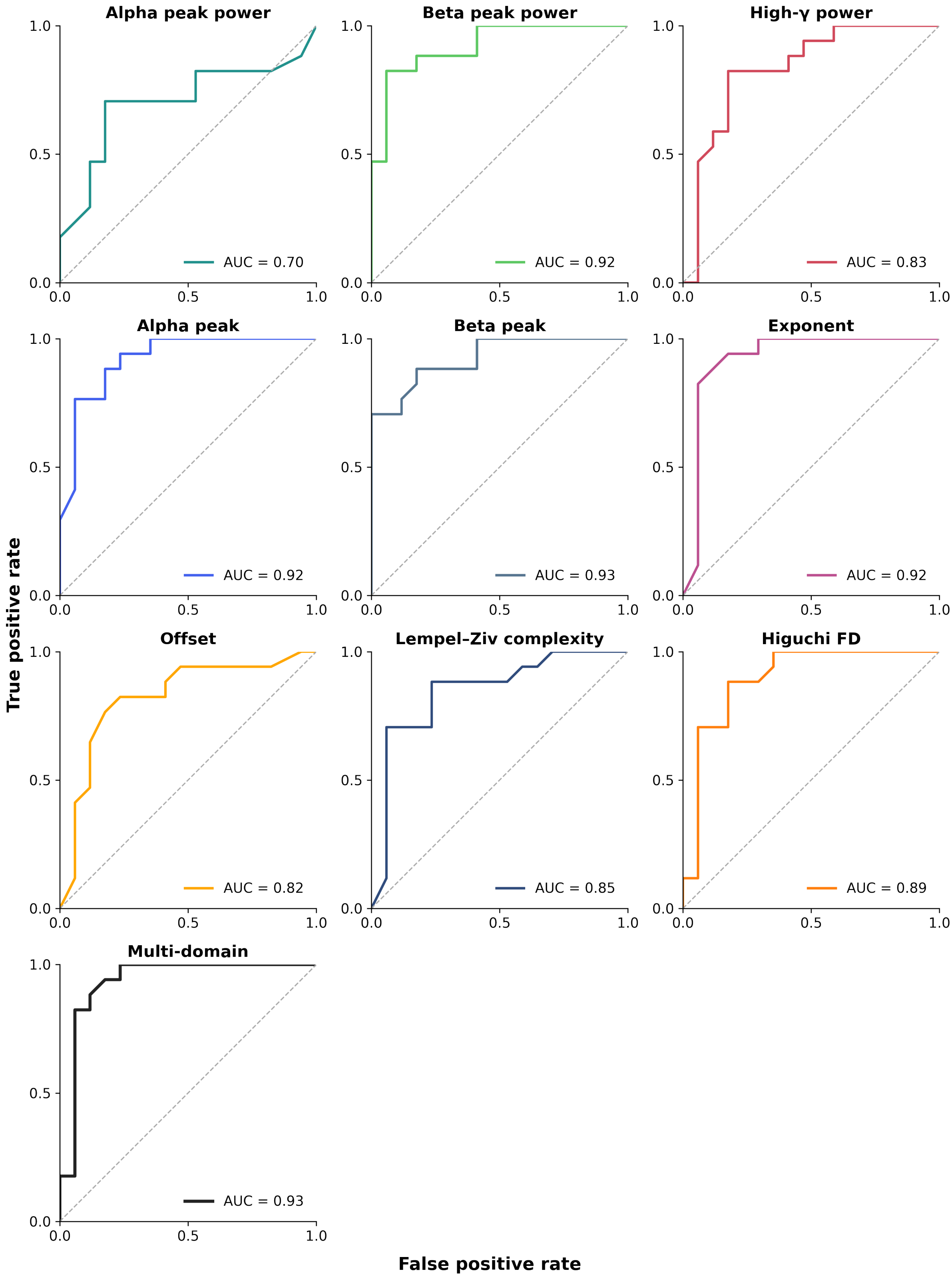


Figure S9: Receiver Operating Characteristic curve of the single-domain and multi-domain ML models. Y axis represents sensitivity and X the false-alarm (1 - Specificity)
